# Multiplexed and reproducible high content screening of live and fixed cells using the Dye Drop method

**DOI:** 10.1101/2021.08.27.457854

**Authors:** Caitlin E. Mills, Kartik Subramanian, Marc Hafner, Mario Niepel, Luca Gerosa, Mirra Chung, Chiara Victor, Ben Gaudio, Clarence Yapp, Ajit J. Nirmal, Nicholas Clark, Peter K. Sorger

## Abstract

High-throughput measurement of cells perturbed using libraries of small molecules, gene knockouts, or different microenvironmental factors is a key step in functional genomics and pre-clinical drug discovery. However, it remains difficult to perform accurate single-cell assays in 384-well plates, limiting many studies to well-average measurements (e.g. CellTiter-Glo®). Here we describe a public domain “Dye Drop” method that uses sequential density displacement and microscopy to perform multi-step assays on living cells. We use Dye Drop cell viability and DNA replication assays followed by immunofluorescence imaging to collect single-cell dose-response data for 67 investigational and clinical-grade small molecules in 58 breast cancer cell lines. By separating the cytostatic and cytotoxic effects of drugs computationally, we uncover unexpected relationships between the two. Dye Drop is rapid, reproducible, customizable, and compatible with manual or automated laboratory equipment. Dye Drop improves the tradeoff between data content and cost, enabling the collection of information-rich perturbagen-response datasets.

## INTRODUCTION

Accurate measurement of cellular responses to perturbation – genetic and drug-induced – is integral to studying regulatory mechanisms and developing new therapies. In the case of small molecules, dose-response studies are increasingly performed at high-throughput using panels of genetically diverse cell lines and compound libraries^1, 2^, with six to nine-point dose-response curves considered the standard for in-depth analysis^3^. When necessary technical and biological repeats are included, a pre-clinical pharmacology profiling study involving ∼100 compounds and ∼50 cell lines can encompass over 10^5^ individual conditions (corresponding to ∼350 384-well plates) – a scale similar to a primary high-content compound screen using diversity libraries or whole-genome screening with RNAi or CRISPR-Cas9 libraries. A key difference between a profiling and screening study is that profiling experiments use a focused set of bioactive compounds or knockouts that target a specific gene family. While most data points contribute to a final profiling dataset, only a small number of hits are typically pursued from chemical diversity or genomic screens. Profiling studies, therefore, benefit greatly from the use of reproducible, sensitive, and relatively low-cost assays.

In any screen, fundamental tradeoffs exist between throughput, number of measurements per condition, cost, and reproducibility; this is true for focused drug and gene panels as well as for genome-scale screens. To increase throughput, cell-based small molecule screens are often performed using single, relatively simple readouts, such as luminescence or well-average ATP levels (measured in lysate)^4^, which are reasonable but far from perfect surrogates for cell viability^5, 6^. For example, mix-and-read assays like CellTiter-Glo® are popular because they are rapid and simple to perform. However, multiplexed assays can extract more information from each condition providing superior insight into mechanism and making follow-up studies more efficient. Multiplexed screening also promises to better identify the cell types and disease states in which a small molecule might have the greatest therapeutic potential^7^. These advances have led to a variety of new, high-content screening methods, which are commonly based on fluorescence microscopy^8^. For example, “cell painting” (five-channel, high resolution, multiplexed imaging of fixed cells) has made single-cell morphological measurements feasible at scale^9,10^. As an alternative to fixed-cell assays, live-cell assays can add detailed information about cell-to-cell heterogeneity and response dynamics^11, 12^. However, live-cell assays remain relatively uncommon in pre-clinical drug discovery because they are perceived to be expensive and require specialized expertise, instrumentation, and data analysis methods.

Any attempt to balance simplicity and cost with information content in a screening or profiling study must consider the substantial up-front expense of maintaining panels of mammalian cell lines and setting up a multi-drug dose-response experiment (personnel, media, multi-well plates, drug treatments, etc.). Thus, methods that extract as much information as possible from each assay are likely economically favorable. Accuracy and reproducibility are also essential^13^. The public release of large-scale drug-response data has been marred by controversy arising from the poor agreement between different databases^14, 15^. We studied the underlying issues^16^ and concluded that much of the problem arises from inherent differences between cell lines that are not adequately accounted for in assay design and data analysis. For example, the failure to consider the impact of cell proliferation rates – which differ between cell lines – on cytotoxic drug response contributes to inconsistency across studies^17^ and obscures relationships between genotype and phenotype. Another common contributor to irreproducibility in high-throughput live-cell or immunofluorescence assays performed in multi-well plates is the uneven loss of cells^18^, particularly cells that are dying or undergoing mitosis (which are less adherent than interphase cells). The extent of cell loss varies with cell type, perturbation, type of assay and operator^16^. Thus, methods performed in multi-well plates are often highly reliable under test conditions yet fail to scale as the conditions become more diverse. Overall, we have found that identifying the precise causes of irreproducibility in a cell screening study can be challenging because the irreproducibility is itself irreproducible^16^.

In search of a simple and economical screening approach that would be robust under diverse assay conditions, we found that a range of multi-step procedures could be performed on live and fixed cells by using a sequence of solutions, each made slightly denser than the last by the addition of iodixanol (OptiPrep™), an inert liquid used in radiology. In this approach, multi-channel pipettes or simple robots add each solution in the series along the edges of the wells in a multi-well plate. This dense solution “drops” gently to the bottom of the well, displacing the previous solution with high efficiency and minimal mixing (testing the method with dyes yielded the “Dye Drop” moniker). This method effectively eliminates the need for mix and wash steps^19^. However, as a practical matter, we found that conventional washing can be performed once live-cell assays are complete and cells are fixed. Thus, the Dye Drop density-based methods can be combined with conventional methods in most cases. Additionally, the Dye Drop procedure helps keep reagent costs to a minimum because the volume needed for each step is lower than with conventional procedures.

In this paper, we describe the development, testing and use of minimally disruptive, customizable, microscopy-based “Dye Drop” and “Deep Dye Drop” assays that use sequential density displacement to collect multiplexed data at low cost and with high accuracy. We describe several ways to implement Dye Drop assays to obtain detailed cell cycle information and quantify single-cell phenotypes that are obscured by population-averages (e.g., the rate of DNA replication or formation of DNA repair foci). Dye Drop methods are an ideal complement to the growth-rate (GR) inhibition method of computing dose responses^16, 20, 21^, and when combined, greatly improve the depth and accuracy of data. They can also be used as an entry point for high-plex immunofluorescence, by CyCIF for example^22^. We also expand on the GR computational framework to make it possible to distinguish cytotoxic and cytostatic drug effects based on Dye Drop data. By collecting a dataset of ∼4,000 nine-point dose-response curves from 58 breast cancer cell lines and 67 small molecule drugs, we demonstrate unexpected diversity in cell division rates and cell cycle distributions under basal and drug-induced conditions. We also show that the cytotoxic and cytostatic drug effects have unexpected relationships to each other and to dose: with some drugs, dosage affects only the fraction of cells arrested whereas with others, the extent of killing varies, and with yet other drugs, both phenotypes vary with dose. Together, these data provide new pharmacological insight and allow us to validate a pipeline of public domain methods and open-source software for performing high throughput, multiplexed dose-response and screening studies at single-cell resolution.

## RESULTS

### Dye Drop assays provide accurate measurements of cell viability in multi-well plates

Errors and irreproducibility in experiments involving adherent cells grown in multi-well plates are thought to have five primary causes: (i) patterned (systematic) biases that arise from edge effects and unequal local growth conditions across a plate; (ii) disturbance and loss of some, but not all, cells in a well due to differences in their properties, notably adhesion; (iii) incomplete exchange of reagents during washing steps due to the use of small volumes and high surface tension; (iv) inconsistent or incorrect data processing; and (v) operator-induced effects arising from differences in how samples are handled (i.e. how reagents are added and cells are washed in multi-well plates)^16, 23, 24^. These factors often interact; for example, high flow rate or agitation during wash or media-exchange steps disturbs dying, mitotic and weakly adherent cells, whereas gentle methods can result in insufficient liquid exchange. Several of these problems become substantially worse as wells become smaller since liquid volumes decrease and surface tension plays a greater role (e.g. 384 vs 96 well plates)^25^. We, and others, have previously described how systematic bias can be mitigated through sample randomization, use of humidified secondary containers etc.^16, 26^.

In this work, we focus on errors introduced by cell loss and uneven reagent exchange during multi-parameter 384-well plate assays on live and fixed cells. Specifically, we sought to develop an approach to reliably assay living cells in multi-well plates at single-cell resolution by optimizing the following factors: (i) simplicity and use of common or commercially available reagents; (ii) minimal disturbance and good retention of delicate and poorly adhered cells; (iii) compatibility with live-cell assays that measure viability, DNA incorporation, and cell cycle progression; (iv) simple customization to enable measurement of proteins and phenotypes relevant to a specific biological problem under investigation; (v) compatibility with simple robots and manual multichannel pipettes; and (vi) cost efficiency, through reduced assay volumes and use of public-domain protocols and software. Preliminary studies established that many existing cell culture assays could be performed in the presence of relatively high concentrations of the density reagent iodixanol^27^. This reagent makes it possible to perform multi-step procedures with a series of increasingly dense solutions (each made denser than the last by addition of increasing iodixanol concentrations). Successive solutions flow to the bottoms of wells and displace the previous solution without aspiration, mixing or disturbing fragile cells. We found that the density displacement (Dye Drop) method is easy to perform without significant training and is compatible with small volumes of solution, reducing the costs of reagents by ∼50% (see below for a detailed discussion).

We next sought to evaluate the accuracy of the Dye Drop method and verify that it does not introduce additional artifacts. To do this, we measured drug-induced growth arrest and cell death using live cell microscopy followed by either a standard wash and fixation or fixation with an iodixanol-containing solution. We used live-cell microscopy of MCF 10A cells expressing the nuclear marker, H2B-mCherry to monitor proliferation and death at single-cell resolution in a time-resolved manner without any fixation or wash steps. This nuclear marker served as a control for the evaluation of Dye Drop methods. MCF 10A cells were exposed for 24 h to one of four cytotoxic drugs (dinaciclib, paclitaxel, staurosporine, or vincristine) at nine doses spanning four orders of magnitude. The YOYO-1 vital dye was added to the medium to detect dead and dying cells, and imaging was performed on a microscope with an environmental control chamber equipped to handle microtiter plates. When live-cell acquisition was complete, cells were fixed with 4% formaldehyde in 10% iodixanol solution in PBS followed by aspiration, addition of PBS to the now fixed cells, followed by another round of imaging. We found that dose-response curves were indistinguishable before and after iodixanol fixation (using cell viability as a measure of response) for each drug recorded (Fig. 1a). In contrast, when live cell imaging was followed by conventional and relatively vigorous washes using an automated plate washer and then by fixation in formaldehyde, we found that the cell number per well decreased following each wash (Supplementary Fig. 1a-b) and that the magnitude of the effect was drug dependent (we explore this in greater detail below).

**Figure 1:**
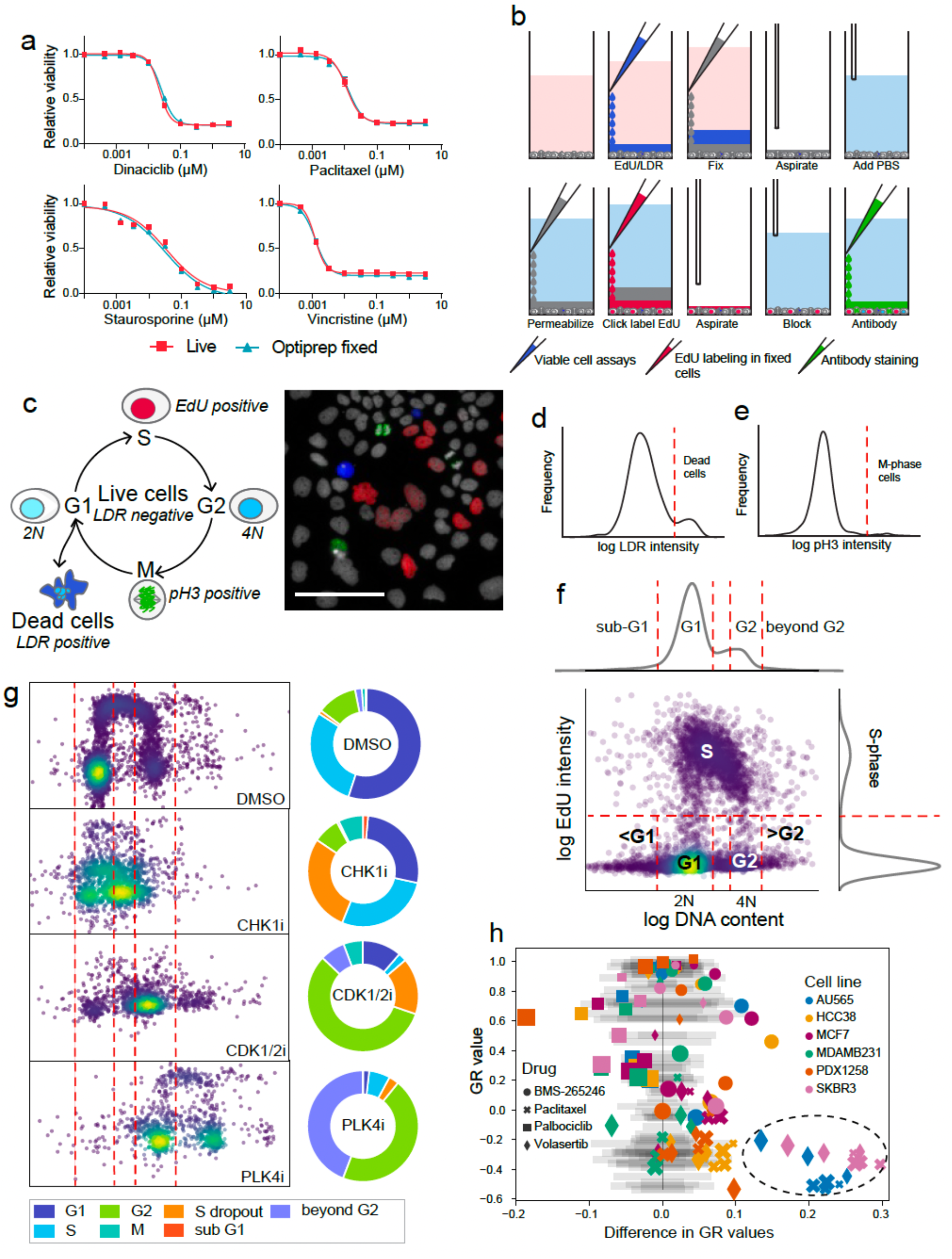
Sequential density-based staining and fixation prevent cell loss from multi-well plates. (a) Relative cell viability in OptiPrep™ fixed cells as compared to live cell microscopy following 24 h treatments with increasing concentrations of dinaciclib, paclitaxel, staurosporine, and vincristine in MCF 10A-H2B-mCherry cells. Error bars represent the standard error of the mean of eight technical replicates from one representative biological replicate. (b) Deep Dye Drop protocol steps: EdU and LDR dye are added in 10% OptiPrep™ followed by fixation with 4% formaldehyde in 20% Optiprep™. Cells are then permeabilized with 0.5% Triton X-100 in 10% OptiPrep, and the EdU is labeled with a fluorescent dye azide via Click chemistry in 20% OptiPrep™. The contents of the well are aspirated, cells are blocked, and then stained with a conjugated antibody against phospho-histone H3 (pH3) in 10% OptiPrep™. One well of a multi-well plate is depicted. (c) Schematic and representative image of cells stained with the Deep Dye Drop protocol. Nuclei are stained with Hoechst (gray-scale), dead cells are stained with LIVE/DEAD red (blue), S-phase cells are labeled with EdU (red) and M-phase cells are stained with phospho-histone H3 (green). Scale bar is 100 µm. (d) Thresholds set to classify dead cells shown on a distribution of LDR intensity values and (e) to identify cells in M-phase shown on a distribution of pH3 intensity values. (f) Scatter plot of EdU intensity versus DNA content. The red dotted lines represent gating applied to assign cells to the sub G1, G1, G2, beyond G2, and S-phases of the cell cycle (see online Methods). (g) DNA content in BT20 cells treated for 72 hours with inhibitors targeting CHK1 (1 µM LY2606368), CDK1/2 (3.16 µM BMS-265246) and PLK4 (0.316 µM CFI-400495) and untreated controls. All cells from a single well in a 384 well plate are shown per condition. (h) The difference in GR values calculated from Deep Dye Drop and conventional assays with respect to the GR value from the Deep Dye Drop assay. The grey bars represent the 95% confidence intervals for GR values from Deep Dye Drop experiments performed in biological triplicate.

### Deep Dye Drop assays enable multiplexed cell viability and cell cycle measurements

We next applied Dye Drop to study the effects of chemical or genetic perturbation on cell viability and cell cycle metrics, such as the rate of division, DNA replication, arrest at discrete cell cycle stages, induction of polyploidy, etc. To do this, we explored whether a Deep Dye Drop could combine live-cell LIVE/DEAD and EdU incorporation assays followed by fixation and processing for immunofluorescence. First, we established that iodixanol did not affect cell proliferation at concentrations needed for this procedure by adding it to cells at concentrations up to 25% for one hour (Supplementary Fig. 1d) or at concentrations up to 5% for 24 h (Supplementary Fig. 1e). Neither a pulsed exposure to a high concentration of iodixanol nor prolonged exposure to a lower concentration had any detectable effect on cell number, consistent with literature describing the use of iodixanol in density gradient purification of viable cells^28^. Based on these data, we then combined the amine-reactive and fixable LIVE/DEAD dye (LDR) with Hoechst 33342^29^ by suspending both in 10% iodixanol in PBS and then adding the solution to wells with a multichannel pipette, thereby displacing the overlying growth medium. Following a 30 min incubation, a solution containing 20% iodixanol and 4% formaldehyde was used to displace the LDR and Hoechst dyes and fix the cells (see online Methods for details; Supplementary Fig. 1f-g). Once fixed, live and dead cells could be easily distinguished by imaging and even dead – potentially weakly adhered cells – were found to be resistant to washing, allowing a variety of staining protocols to be followed, as described below.

To monitor DNA replication, EdU was added to cells at the same time as the LDR viability dye (but without Hoechst 33342), resulting in its incorporation by cells actively synthesizing DNA. Following fixation, EdU incorporated into DNA was fluorescently labeled using Click chemistry to visualize S-phase cells. M-phase cells were then stained with an anti-phospho-histone H3 antibody (anti-pH3; which is available covalently coupled to fluorophores, thereby avoiding the need for secondary antibodies) (Fig. 1c-f). Incubating cells in the presence of antibody overnight resulted in good quality staining and provided a natural breakpoint in the protocol; Hoechst 33342 staining was performed in parallel. Imaging viable cells processed this way generated the classic “horseshoe” profile of DNA synthesis and content^30^, enabling detailed analysis of DNA replication rates and S-phase errors; it also reliably discriminated G1 and G2 populations and detected cells with aberrant DNA content (Fig. 1g-h, Supplementary Fig. 1h). Moreover, the now fixed plates could be subjected to additional staining protocols with the option of using either additional Dye Drop steps to economize on reagents or switching to conventional plate washing methods.

To compare the resulting Deep Dye Drop assay to equivalent multiplexed staining achieved by conventional assays under real-world conditions, we optimized the settings on an Agilent BioTek EL406 plate washer so they would be as gentle as possible while also ensuring effective reagent exchange. The EL406 instrument is prototypical of compact multi-well plate washing robots found in many academic and industry screening facilities; this instrument is also inexpensive enough for a single lab to purchase. We then exposed six widely used breast cancer cell lines (a subset of the lines described below) to nine-concentration of each of four drugs having different mechanisms of action: CDK1/2 inhibitor BMS-265246, CDK4/6 inhibitor palbociclib, microtubule stabilizing drug paclitaxel, and PLK1 kinase inhibitor volasertib. These drugs arrest cells at distinct points in the cell cycle, and in many lines also induce cell death. Parallel assay plates were processed using Deep Dye Drop or a standard high-throughput protocol (see Methods) and responses quantified using GR values. We then plotted the absolute GR for each data point (as measured by Deep Dye Drop) against the observed difference in GR value obtained by Deep Dye Drop and standard assays across all drugs (Fig. 1b; shapes), cell lines (colors) and doses (symbol size; the underlying grey bars represent the 95% confidence interval for triplicate Deep Dye Drop experiments as a means of comparison). These data revealed cell-line and drug dependent differences in GR measurements between the two approaches with the greatest differences observed in the cases of AU565 and SKBR3 cells treated with paclitaxel or volasertib, which are conditions that induce substantial cell killing (see also Supplementary Fig. 1c). These data are consistent with previous results showing that condition-dependent effects on cells are a primary contributor to data irreproducibility.^16^ Thus, in comparison to a conventional staining techniques, the Deep Dye Drop approach improves accuracy and better preserves cells that are vulnerable to loss.

We then attempted to estimate the costs of CellTiter-Glo®, conventional staining, and Dye Drop protocols based on an exemplary study performed by an experienced technician in Boston, MA in 2022 involving 12 cell lines (processed in two batches of 6 cell lines each) and 30 drugs, with each drug assayed in triplicate at nine concentrations per replicate (Supplementary Fig. 1i, **Supplementary Table 1**). The bulk cost for this type of study is the time and materials required to grow multiple cell lines and seed multiple 384-well plates – 48 plates were needed for this experiment. This expense was the same for all protocols ($170 per plate). Reagent costs for CellTiter-Glo® assays were approximately two-fold higher ($52 per plate) than for microscopy-based Deep Dye Drop assays ($28 per plate for assaying viability, EdU incorporation and one immunofluorescent marker) and 10-fold higher than for simpler Dye Drop assays ($5 per plate for viability alone). Were one to mimic the full set of Deep Dye Drop assays using conventional staining and fixation procedures (ignoring for the moment problems with cell displacement and reagent exchange), reagent costs would be about three-fold higher than with Deep Dye Drop due to larger working volumes per well. However, in comparing CellTiter-Glo® with imaging-based assays we must also account for the fact that plate scanning microscopes require more expensive plates than luminescent plate readers and more time to perform (increasing salary costs). Accounting for all these factors and including initial tissue culture, we found that the final assay steps (Deep Dye Drop or CellTiter-Glo®) represented only about 10% of the overall cost (**Supplementary Table 1**). Thus, multiplexed single-cell assays can be performed at scale at roughly the same cost as a well-average CellTiter-Glo® measurement while extracting vastly more information on cell viability, cell cycle state, DNA incorporation and one or two marker proteins. We conclude that, in many settings, Deep Dye Drop assays are likely to be the preferred way to measure cell perturbations.

### Customizing Dye Drop assays for different endpoints

To customize the Deep Dye Drop method for different biological endpoints, we tested a range of antibodies (Fig. 2a-e) and found that it was straightforward to vary the immunofluorescence component of the Deep Dye Drop protocol for additional biological insight. For example, we treated MCF7 cells with the CDK4/6 inhibitor palbociclib and assayed them with Deep Dye Drop using different antibodies. We first used an antibody against phospho-pRb, which allowed us to measure dose-dependent target engagement at a single cell level (i.e. drug pharmacodynamics; Fig. 2a) and the degree of G1 arrest. We also stained palbociclib-treated MCF7 cells with an anti-beta-actin antibody and detected a change in cell shape upon drug treatment (Fig. 2b)^31, 32^. Similarly, treating MCF 10A cells with the topoisomerase II inhibitor, etoposide and staining them with an anti-53BP1 antibody, revealed that the fraction of cells with multiple 53BP1-containing DNA damage foci increased in a dose-dependent manner (Fig. 2c). In this case, we used an anti-mouse secondary antibody to show that the addition of immunofluorescence to Deep Dye Drop does not require fluorophore-conjugated antibodies. Finally, we stained actinomycin D-treated MCF7 cells with cytochrome C and were able to quantify mitochondrial outer membrane permeabilization (MOMP; a key step in the commitment to apoptosis^33^) based on changes in cytochrome C localization (Fig. 2d). Thus, given a suitable antibody for immunofluorescence, Deep Dye Drop assays can be used to measure many molecular processes at a single cell level in normally growing and perturbed cells. Of note, phenotypes such as cell flattening, DNA focus formation, and MOMP are not readily detectable using well-average measurements (ELISA assay for example) or multiplexed flow cytometry.

**Figure 2:**
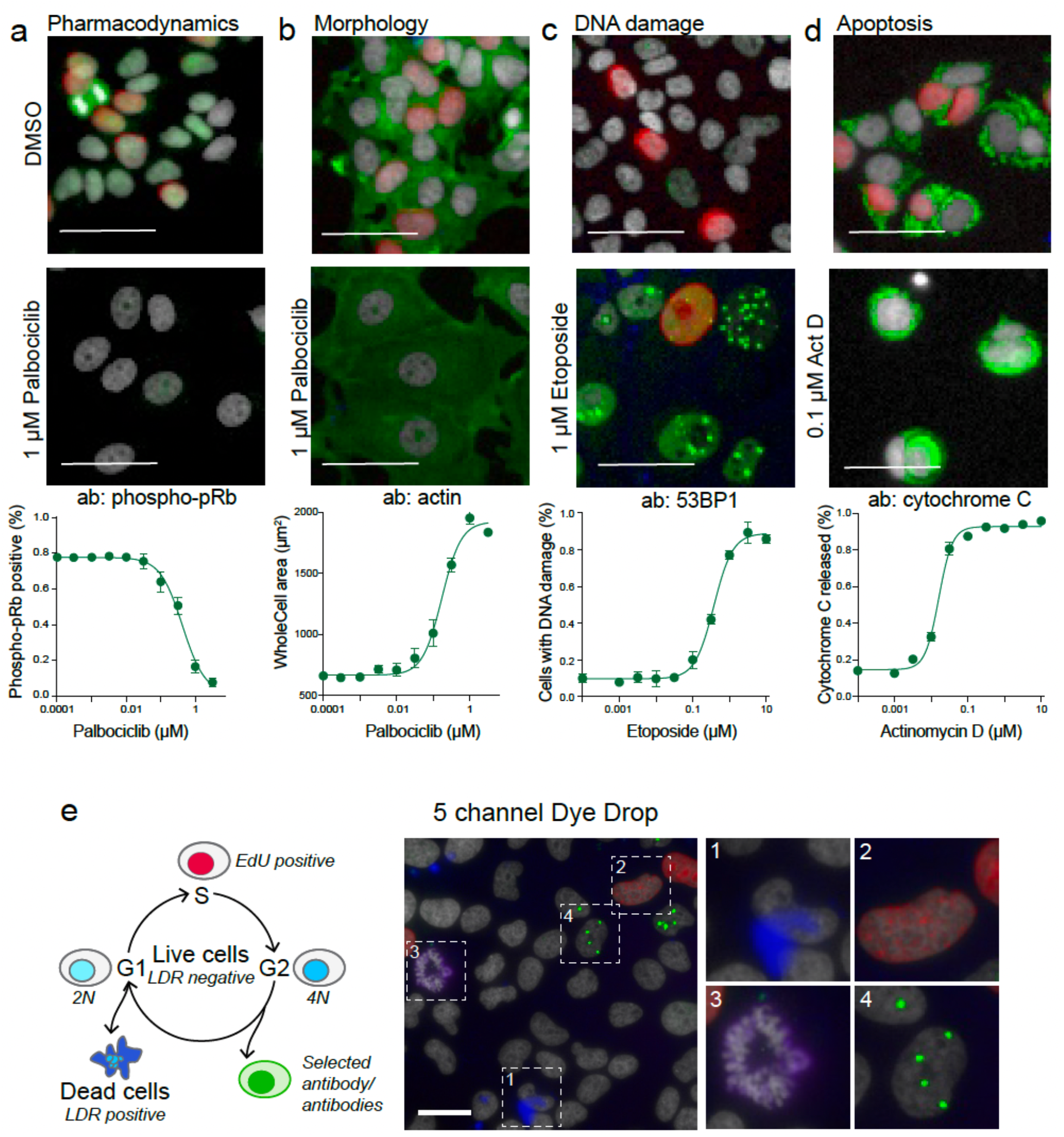
Customization of antibody incorporation in Dye Drop assays. (a) MCF7 cells stained with phospho-pRb and (b) actin untreated and after 72 h in 1 µM palbociclib; effects of increasing concentrations of palbociclib on the fraction of phospho-pRb positive MCF7 cells, or (b) on cell size as detected with actin staining after 72 h. (c) MCF 10A cells stained with 53BP1 untreated and after 72 h in 1 µM etoposide; induction of DNA damage by increasing concentrations of etoposide in MCF 10A cells as detected with 53BP1 staining after 72 h. (d) MCF7 cells stained with cytochrome C untreated and after 72 h in 0.1 µM actinomycin D; effect of increasing concentrations of actinomycin D on release of cytochrome C from the mitochondria in MCF7 cells after 72 h, performed in duplicate. Nuclei are stained with Hoechst (gray-scale), and EdU (red) in all images. Error bars represent the standard deviation of the mean of four replicates. Scale bars are 50 µm. (e) Schematic and representative image of the addition of a fifth channel to the Deep Dye Drop assay: cells are stained with Hoechst (gray-scale), LDR (blue, 1), EdU (red, 2), pH3 (purple, 3) and 53BP1 (green, 4). Scale bar is 50 µm.

Most modern fluorescence microscopes are equipped to measure five or more fluorescent channels. To develop a five-channel Deep Dye Drop protocol, we performed dual antibody staining and identified five complementary and commercially available assays that could readily and reproducibly be performed in a 384 well format: (i) LIVE/DEAD assays, (ii) live-cell EdU incorporation, (iii) fixed cell counting with DNA content and morphology in the Hoechst channel, and (iv-v) two-channel immunofluorescence using Alexa 488 and Alexa 750-conjugated primary or secondary antibodies (anti-pH3 and anti-53BP1 primary antibodies were used in Fig. 2e, but antibody selection should be adapted to the biological questions being pursued).

Several methods have recently become available to collect multiplexed immunofluorescence data from cells grown in culture^34^ as a means to obtain detailed insight into single cell states; such methods are a logical follow-on to Deep Dye Drop. Cyclic immunofluorescence (CyCIF)^35^, for example, is a public domain method that enables collection of 10-20 plex images through sequential rounds of 3 or 4-plex antibody staining, imaging, and fluorophore oxidation (Fig. 3a). We found CyCIF and Deep Dye Drop assays to be compatible with only slight modification – it was necessary to use EDTA to quench the copper in the click chemistry used for EdU labeling prior to adding the hydrogen peroxide-containing fluorophore inactivation solution used for CyCIF. Alternatively, the click reaction could be performed after all CyCIF staining cycles were complete. We also found that it was possible to introduce a gap of up to several weeks (after plates were fixed) between an initial Deep Dye Drop assay and CyCIF. This enables Deep Dye Drop analysis to inform the choice of antibodies for CyCIF and help focus the more complex and expensive assays on a subset of informative conditions.

**Figure 3:**
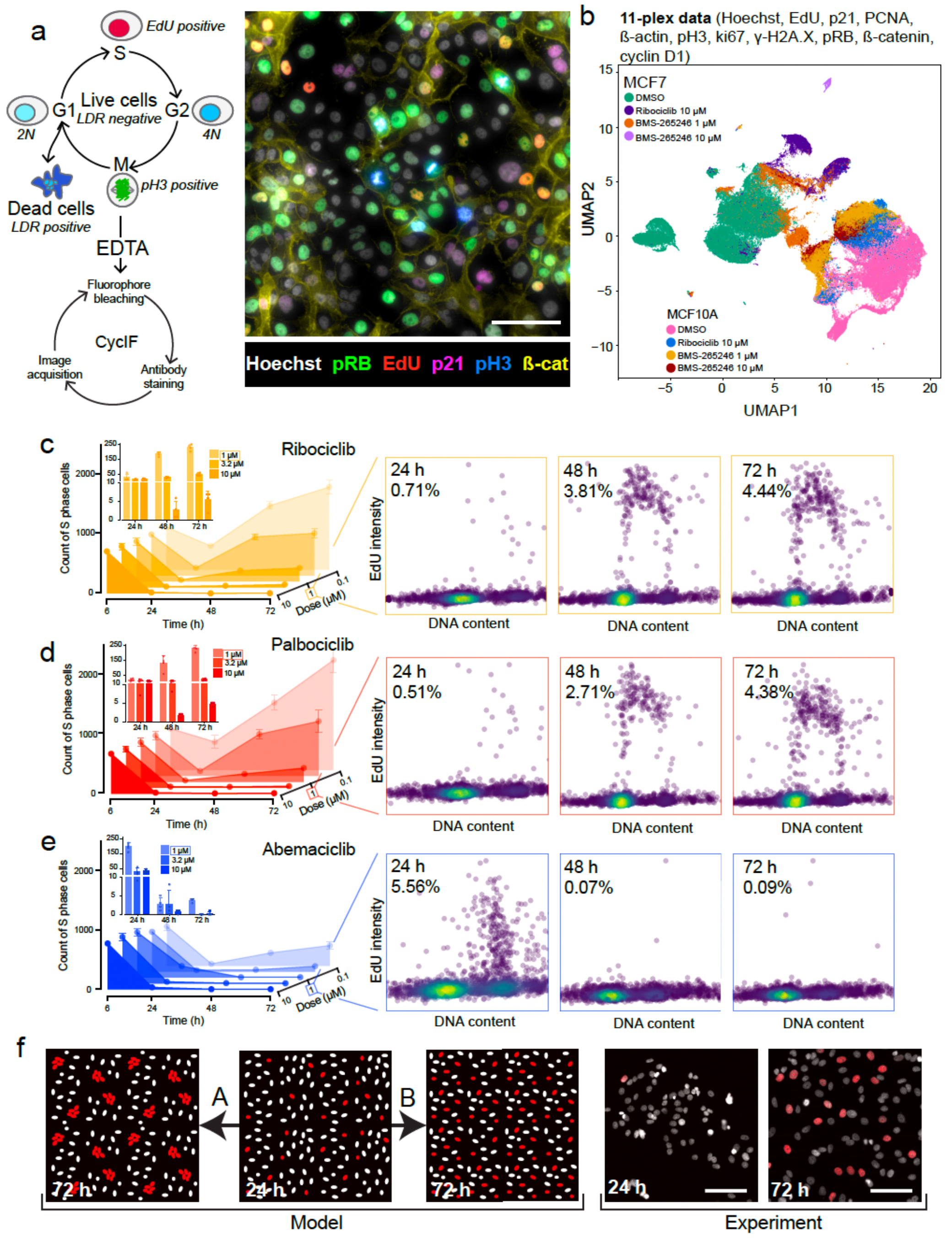
Integration of Dye Drop assays with CyCIF and time series. (a) Schematic and representative image of multiplexing Deep Dye Drop assays with cyclic immunofluorescence: the Hoechst (gray-scale), EdU (red), pH3 (green), beta-catenin (cyan), phospho-pRb (blue), and p21 (yellow) signals are displayed, and contrast was adjusted for visualization purposes only. Scale bar is 100 µm. (b) UMAP representation of MCF7 and MCF 10A cells treated with BMS-265246 (1 µM, 10 µM), ribociclib (10 µM) or DMSO stained with Deep Dye Drop and cyclic immunofluorescence. (c) The number of MCF7 cells in S-phase following treatment with increasing concentrations of ribociclib, (d) palbociclib, and (e) abemaciclib after 6, 24, 48, and 72 h. EdU versus DNA content scatter plots show the single cell cell-cycle distribution at 1 µM doses of each drug at the time points indicated, the percentage of cells in S-phase is indicated on each plot. Error bars represent the standard deviation of four technical replicates. The scatter plots show all cells in a single, representative well of a 384 well plate for each condition. (f) Illustration of possible patterns of the emergence of resistant cells. A, clonal, genetic resistance or B, non-clonal, non-genetic adaptation followed by images of MCF7 cells treated with 1 µM palbociclib for 24 h or 72 h. Nuclei are visualized with Hoechst in white and EdU positive cells are shown in red, scale bars are 100 µm.

To illustrate the integration of Dye Drop assays with CyCIF, we treated MCF7 and MCF 10A cells with ribociclib (CDK4/6 inhibitor that induces G1 arrest), BMS-265246 (CDK1/2 inhibitor that induces G2 arrest and toxicity) or DMSO for 72 h, then performed Deep Dye Drop staining followed by three rounds of CyCIF staining (an 11-plex measurement). When the multiplexed single cell CyCIF data were visualized using UMAP (Uniform Manifold Approximation and Projection for Dimension Reduction)^36^, we found that MCF7 and MCF 10A cells clustered separately, as did cells treated with each drug (Fig. 3b **and** Supplementary Figure 2a-d; note that 10 µM BMS-265246 was cytotoxic; therefore, the UMAP projection contains fewer cells for that condition). What is striking about these data is that both CDK1/2 and CDK4/6 inhibitors generated two different arrest states that are not the same as those traversed by normally dividing cells. These states are not distinguished by any single marker in our antibody panel, but rather are by high dimensional features. This complexity in cell cycle arrest states is consistent with recent evidence from human tumors^37^ and reveals how CyCIF can be used to discriminate drug-induced cell states that appear similar by lower-plex assays. It seems likely that such data, when collected at scale, will assist in identifying drug mechanism of action and response biomarkers.

Acquired and adaptive resistance to therapy is a barrier to successful cancer treatment and an area of intense focus in pre-clinical research.^38^ To illustrate the use of Dye Drop assays in studying this phenomenon, we exposed hormone receptor positive (HR^+^) MCF7 cells to three related CDK4/6 inhibitors approved to treat HR^+^/HER2^-^ breast cancer (palbociclib, ribociclib, and abemaciclib)^39, 40^. We found that cell cycle was effectively inhibited, with cells accumulating in G1, within 24 hours of drug exposure but that only abemaciclib was able to sustain G1 arrest; in cells exposed to palbociclib and ribociclib, the fraction of S-phase cells increased 5 to 8-fold between 24 and 72 h (Fig. 3c-e). The greater efficacy of abemaciclib is likely due to its inhibition of multiple CDKs in addition to CDK4 and CDK6^41^ (Fig. 3c-e and inset plots). To study the frequency of cell cycle re-entry at a single cell level, we asked whether MCF7 cells that had started to grow in the presence of palbociclib or ribociclib were found in clusters (colonies) or were distributed across the dish. Acquired drug resistance generally involves the outgrowth of clones arising from a low-frequency mutation. In contrast, adaptive drug resistance involves a non-genetic adaptation arising in many cells in a population (Fig. 3f)^42^. Following 24 h of 1 or 10 µM palbociclib exposure, we found that the physical distance between S phase cells increased (Supplementary Fig. 2e-f) consistent with a reduction in their number and arrest in G1 (Fig. 3d, f). By 72 h, cells had started to adapt to the drug and S-phase fraction increased ∼8-fold but these cells were as far apart as at 24 hr. Thus, drug resistance arose frequently throughout the population and not in clones, consistent with rapid adaptation rather than rare mutation^43^. These data illustrate the ability of spatially-resolved single-cell data, as opposed to well-average measurements, to provide valuable insight into drug resistance.

### Using Dye Drop to systematically screen small molecule drugs in breast cancer cell lines

To demonstrate the use of Dye Drop assays at scale, a panel of 58 breast cancer cell lines was exposed to a library of 67 approved and investigational kinase inhibitors and other small molecules. The panel included multiple non-malignant breast epithelial lines (labeled NM) and many cell lines routinely used to study the three major breast cancer subtypes: hormone receptor positive (HR^+^), HER2 amplified or overexpressing (HER2^amp^), and triple negative (TNBC; which lack expression of estrogen, progesterone and HER2). Responses were measured 72 h after the addition of drug at nine concentrations spanning a 10^4^ dose range (plus DMSO-only negative controls). All assays were performed in triplicate or quadruplicate to measure technical repeatability. This yielded a total of ∼3,900 nine-point dose response curves computed from ∼116,000 wells (∼35,000 unique conditions), with each well yielding data on ∼0.5 to 15 x10^3^ single cells (Fig. 4a, Supplementary Fig. 3).

**Figure 4:**
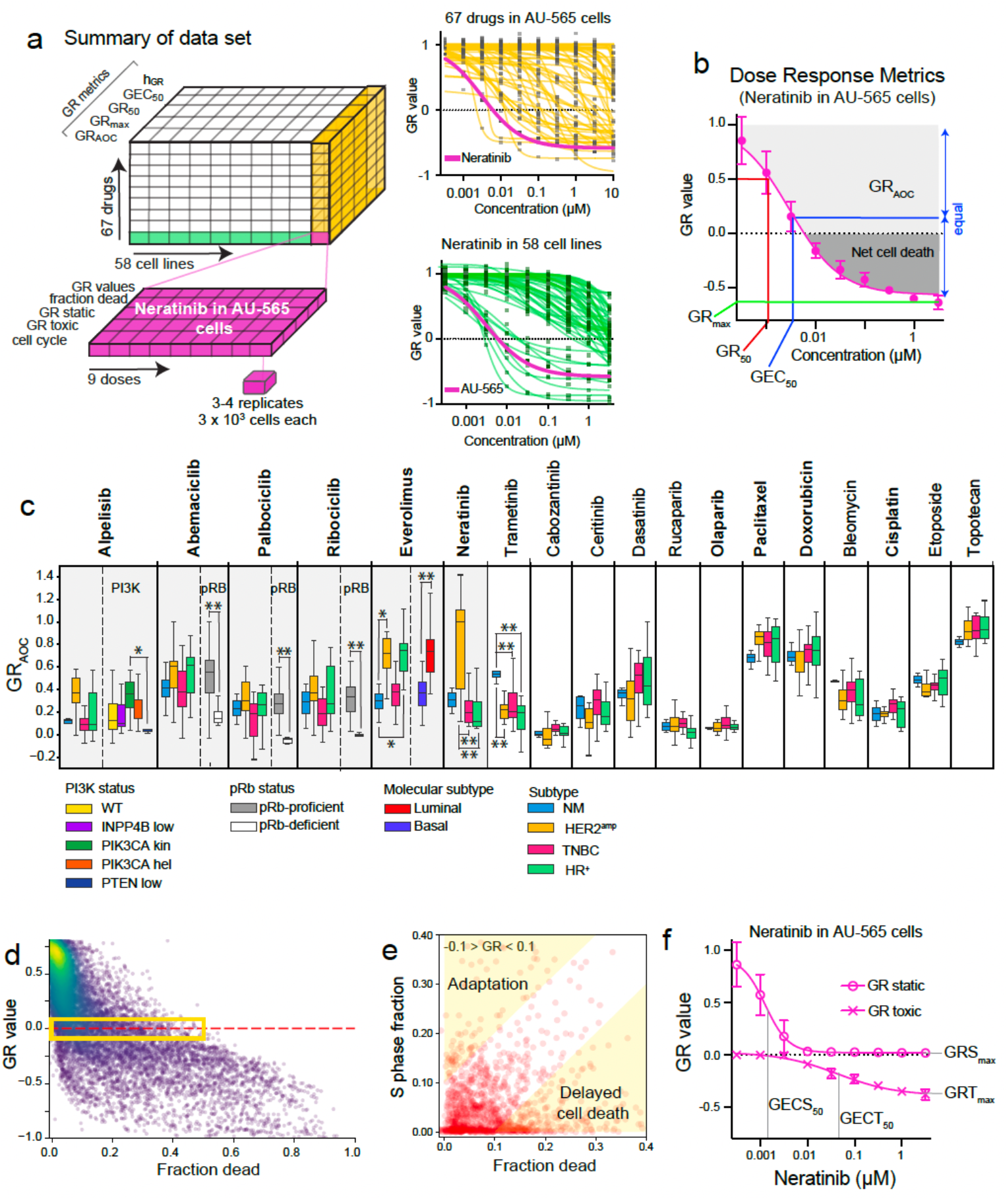
Dye Drop screening of a breast cancer cell line panel treated with a kinase inhibitor library. (a) Dye Drop and Deep Dye Drop assays were used to measure the responses of 58 breast cancer cell lines treated with 67 drugs at nine concentrations after 72 h in technical triplicate or quadruplicate. GR-dose response curves for all drugs in AU-565 cells and all cell lines treated with neratinib. The response of AU-565 cells to neratinib is shown in pink. Error bars are omitted for simplicity. (b) Illustration of GR metrics shown on the neratinib-AU-565 dose response curve. Error bars represent the standard deviation of technical quadruplicates. (c) Boxplot of GR_AOC_ values for FDA-approved drugs included in our screen. Cell lines are separated by clinical subtype (n = 5 NM, 13 HER2^amp^, 13 HR^+^, 26 TNBC), dose responses were measured in technical triplicate or quadruplicate. The bottom and top of the box show the first and third quartiles, the bar within each box represents the median value and error bars represent the range of values. * P-value < 0.05, ** P-value < 0.01, 2-way ANOVA with Tukey’s correction for multiple comparisons, or unpaired 2-tailed t-test when only comparing two groups. The shaded region indicates responses that align with known biomarkers. (d) Variability in fraction dead with respect to GR value for all dose response data, GR values > 0.8 are not shown. (e) Fraction of cells in S phase relative to fraction dead for conditions resulting in −0.1 > GR < 0.1. Regions indicative of late-onset cell death and adaptation to drug are highlighted in yellow. (f) GR^S^ and GR^T^ curves for AU-565 cells treated with neratinib. Error bars represent the standard deviation of technical quadruplicates.

GR values were determined for each drug, dose and cell line based on the number of viable cells at t = 0 h and 72 h followed by curve-fitting to estimate the four primary metrics of drug response^17^: (Fig. 4b): (i) potency, as measured either by GR_50_ (the drug concentration at which GR = 0.5) or by GEC_50_ (the concentration at half-maximum effect), which is relevant when the magnitude of drug-induced arrest or killing is insufficient for GR = 0.5 to be reached; (ii) efficacy, as measured by GR_max_ (the maximum drug effect, typically achieved at the highest dose); (iii) the slope of the fitted dose response curve, h_GR_; and (iv) and the integrated drug effect as measured by the area over the GR curve GR_AOC_ (AOC in this setting is directly analogous to AUC data used with other drug response metrics)^44–46^. Drug response data exhibited good reproducibility (median standard deviation of GR values = 0.07 and median coefficient of variation ∼11%) and recapitulated genetic associations observed in the clinic. For example, PIK3CA-kinase domain mutant lines were significantly more sensitive than PTEN-low^47^ lines to alpelisib (PIQRAY®, approved for PIK3CA-mutant HR+/HER2-advanced and metastatic disease; 2-way ANOVA P-value < 0.05)^48^; HER2^amp^ lines were more sensitive to neratinib (NERLYNX® approved for HER2^amp^ disease; 2-way ANOVA P-value < 0.01)^49^; luminal lines were more sensitive than basal lines to everolimus (AFINITOR® approved for HR^+^ disease; two-tailed unpaired t-test P-values < 0.001)^50^; and pRb-deficient lines^41^ were resistant to the CDK4/6 inhibitors palbociclib, ribociclib and abemaciclib (IBRANCE®, KISQALI®, and VERZENIO®; two-tailed unpaired t-test P-values < 0.001) (Fig. 4c; see Supplementary Fig. 4a for all others). Several additional drugs exhibited subtype specificity that matched their targeted clinical indication. For example, relative to other subtypes, HR^+^ cell lines were more sensitive to the AKT inhibitors AZD5363 and ipatasertib, and to the mTOR inhibitors everolimus, AZD2014 and LY302341; TNBC cell lines were more sensitive to the WEE1 inhibitor adavosertib, the ATR inhibitor AZD6738, and the PARP inhibitors rucaparib and olaparib^51–53^. Observing such associations between clinical and preclinical data is not trivial: we have previously shown that it requires accurate phenotypic measurement and the use of growth rate-corrected drug response metrics^21^.

One striking feature of these data is that non-transformed (NM) cell lines were not on average more resistant to drugs than cancer lines; this was true of pre-clinical compounds and drugs approved by the FDA for treatment of breast cancer (Fig. 4c, in bold). As a whole, NM cells were actually more sensitive than tumor cells to the ERK1/2 inhibitor BVD523, and to the MEK1/2 inhibitor trametinib (two-tailed unpaired t-test, P-value < 0.05). This is not a new observation^54^, but it demonstrates that even with approved therapeutics and a panel of non-transformed and cancer cell lines, we should not expect to observe a consistent difference in drug response between cells representative of normal and diseased states.

### Distinguishing cytostatic and cytotoxic drug effects

To further explore the biological implications of GR values, we looked more closely at the balance between cell birth, death, and arrest in our dataset. A value of GR = 1 corresponds to unperturbed proliferation resulting in identical numbers of viable cells in drug-treated and control cultures; GR = 0 corresponds to no increase in viable cell number; and GR < 0 to net cell loss. In principle, the GR = 0 condition could arise because all cells in a culture arrest in a viable state (true cytostasis) or because the number of cells that die equals the number born during the assay. To distinguish these possibilities, we compared the fraction of dead cells to GR values across all drugs, concentrations, and cell lines (Fig. 4d). We observed the anticipated negative correlation between GR value and dead cell fraction but with high dispersion around the trend line: at GR = 0, the fraction of dead cells varied from 0 to 48% (median = 12%). We then compared the drug-induced change in S-phase and dead cell fractions (for −0.1 < GR < 0.1) and found that birth and death balanced (S-phase and death fractions within 5% of each other) fully in ∼44% of conditions, resulting in the absence of net cell growth. In a further ∼24% of conditions, high cell death was accompanied by low S-phase fraction (i.e. when the difference was >10%); these are likely cases in which GR = 0 is a transient phenomenon preceding cell death (many anti-cancer drugs exert their cytotoxic effects after several days of delay)^55, 56^. The opposite scenario of low cell death with high S-phase fraction (observed in ∼7% of conditions) may reflect adaptation: maximum drug effect likely occurred at an earlier time point and by 72 h, cells had resumed S-phase (Fig. 4e). In such cases, the underlying assumption in GR calculations that growth rate is constant is violated and time-dependent GR measurements are required (Fig. 3c-e). Based on this analysis, we conclude that true cytostatic cell-cycle arrest is likely to occur in only ∼25% of GR= 0 conditions, or ∼2% of all conditions assayed in total with balanced proliferation and death about twice as common.

We identified two limitations with these data. First, we found that a subset of cultured cells died in the absence of any drug treatment (median value 5%; range ∼1 – 19% depending on cell line). Cytotoxicity was therefore computed as the difference in dead cell count between samples with and without drug present. Second, we found that a subset of cells undergoing programmed cell death lysed completely and were therefore not captured as LDR-positive cells in the “dead cell count.” In the absence of continuous live-cell imaging it is not possible to quantify the fraction of cells undergoing lysis, but we surmised that it varied with condition and was greatest when GR ∼ −1.0 (i.e. under highly cytotoxic conditions) and the total cell count (live + dead) was much lower than the time = 0 cell count. A low fraction of dead cells under cytotoxic conditions can be misleading when the total cell count is low (i.e., when many dead cells have lysed); it is therefore important to flag cases in which both low cell number and low cell death co-occur. These limitations do not weaken our conclusion that true cytostasis is less common than balanced birth and death; if anything, lysis of dead cells leads to an underestimate of the extent of cell killing.

To further distinguish between cytostasis and cytotoxicity under conditions when cells are both dividing and dying, we estimated rates of cell transition from proliferation to stasis (k_s_) or death (k_d_) using an ordinary differential equation (ODE) model of cell proliferation. We then computed dose-response curves and metrics for cytostatic and cytotoxic responses (GR^S^ and GR^T^). Note that, while GR^S^ and GR^T^ values can be compared to each other across conditions, they are based on transition rates and therefore do not sum up to the GR value; instead, GR values are approximately equal to the product of GR^S^ and GR^T^ (see methods for details; Fig. 4f, Supplementary Fig. 4b). With data comprising only two time points, the model is formulated such that k_s_ and k_d_ are constant over the course of the experiment (the collection of time-series data overcomes this limitation). Using neratinib response in the HER2^amp^ AU-565 cell line as a case study, we found that the GR values were well fit by a sigmoidal curve (h_GR_ = 0.84) that exhibited both high potency (GR_50_ = 1.2 nM, GEC_50_ = 3.0 nM) and high efficacy (high cell killing; GR_max_ = −0.70) (Fig. 4a-b). When the response was decomposed, the cytostatic component was 40-fold more potent (GEC^S^_50_ = 1.1 nM) than the cytotoxic component (GEC^T^_50_ = 48 nM). Thus, at low neratinib concentrations, cell cycle arrest predominated, but arrest and death co-existed at drug doses near the serum C_max_ in humans^57^ (Fig. 4h). In support of this conclusion, EdU incorporation exhibited half-maximal inhibition of DNA synthesis at 1.5 nM, consistent with cell cycle arrest at this concentration and a requirement for HER2 in cell cycle progression^58^; in contrast LDR data confirmed half-maximal cell killing at ∼30 nM (Supplementary Fig. 5). In general, we found that cytostatic effects were elicited at lower concentrations than cytotoxic effects although in the case of drugs such as dinaciclib, cytostatic and cytotoxic drug concentrations were very similar (median 2-fold difference in GEC^S^_50_ vs GR^T^_50_ across all cell lines).

Maximal cell killing (GR^T^_max_), and potency (GEC^S^_50_) varied widely across the dataset (GR^T^_max_ = 0.06 to −0.74; GEC^S^_50_ median = 0.49 µM, interquartile range (IQR) = 4.85 µM) and were correlated (Spearman r = 0.41, P-value < 0.001) (Fig. 5a; Supplementary Fig. 4b). However, the overall correlation masked different relationships between dose, cytostasis, and cell killing for different drugs. For example, the potency of palbociclib, ribociclib and the CDK4/6-targeting BSJ-03-124 PROTAC (PROteolysis Targeting Chimera; a drug that induces proteasome-mediated degradation of a target)^59^ varied widely across cell lines (GEC^S^_50_, median = 0.54 µM IQR = 8.2) but induced little or no cell death (GR^T^_max_ ∼ 0). In contrast, varying the concentrations of dinaciclib and alvocidib, which target multiple CDKs (i.e., CDK1/2, CDK4/6, CDK5, 7, 9) changed the fraction of cells killed (GR^T^_max_= 0 to −0.7 for both), but the potency remained nearly constant (GEC^S^_50_ = 15 ± 7 nM for dinaciclib; GEC^S^_50_ = 150 ± 70 nM for alvocidib). Compounds such as abemaciclib^41^ and BSJ-03-123^60, 61^ – which have multiple CDK targets – varied on both potency and efficacy axes. A similar pattern was observed for the CDK7 inhibitor YKL-5-124, which varied in potency and efficacy, whereas drugs with activity against CDK12/13 such as THZ1 (CDK7/12/13) and THZ531 (CDK12/13) varied primarily in cytotoxicity. Thus, deconvolution of Dye Drop dose-response data reveals unique modes of anti-cancer drug action with dose (Fig. 5b). The reasons for these differences, and their relationship to target specificity and activity in patients requires further study. However, we note that drugs in our collection that varied little in potency, such as dinaciclib and alvocidib, failed in trials due to excess toxicity, whereas drugs such as abemaciclib that vary in both potency and efficacy appear to be superior as human therapeutics to drugs that induce little or no cell killing, such as ribociclib^41, 62^.

**Figure 5:**
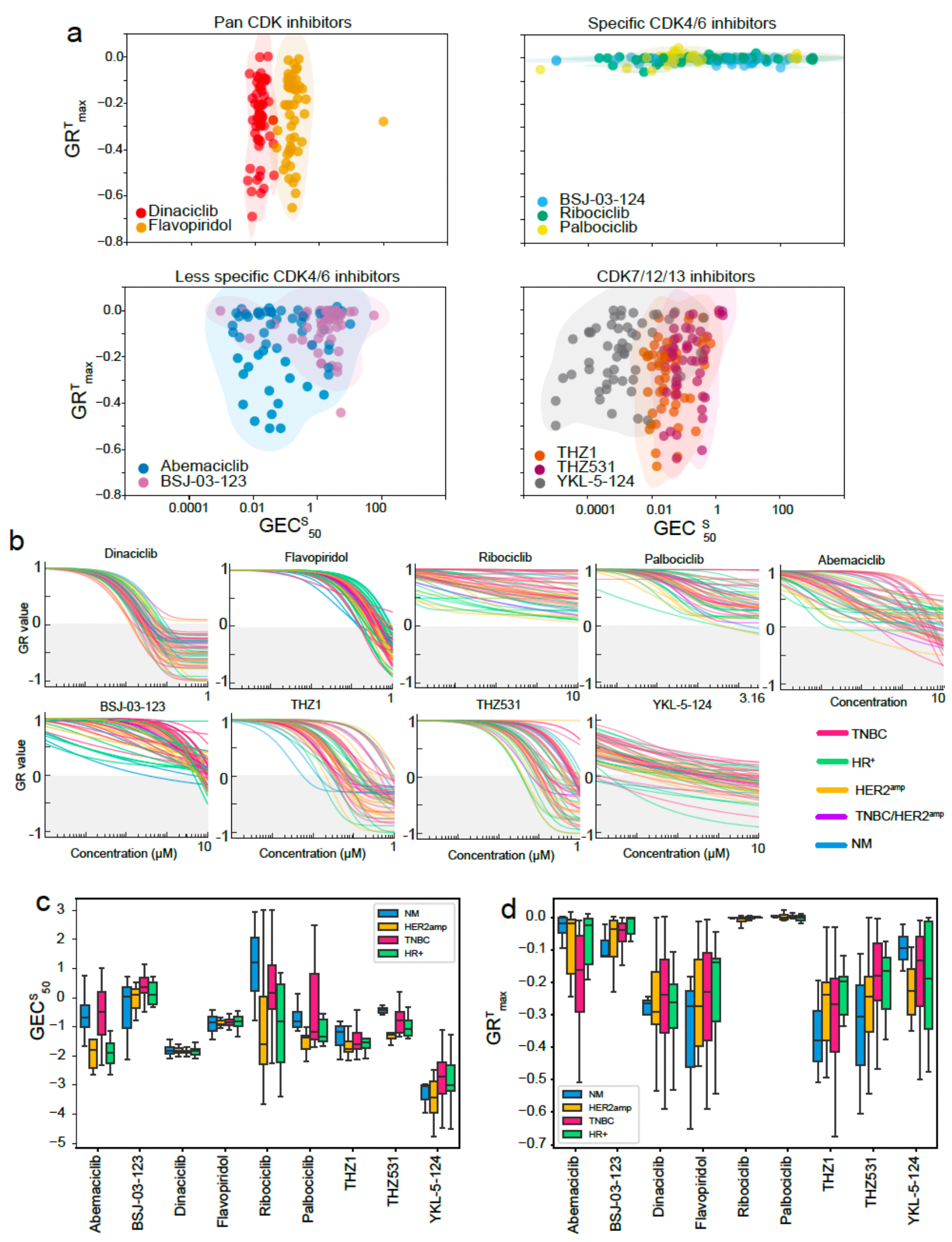
Variation in static and toxic components of responses to CDK inhibitors. (a) Relationship between GR^T^_max_ and GEC^S^_50_ for drugs targeting pan-CDKs (top-left), CDK4/6 (top-right), CDK4/6 and off-targets (bottom-left) and CDK7/12/13 (bottom-right). Shading is for visualization only. (b) The GR dose response curves for the same conditions shown in (a) 72 h after drug addition. Summary boxplots of (c) GECS50 and (d) GRTmax values by subtype for the same conditions shown in (b), the bottom and top of the box show the first and third quartiles, the bar within each box represents the median value and error bars represent the range of values. n = 5 NM, 13 HER2^amp^, 13 HR^+^, 26 TNBC and 1 TNBC/HER2^amp^ cell lines. The TNBC/HER2^amp^ cell line was excluded from (c) and (d) for simplicity. Dose responses were measured in technical triplicate or quadruplicate.

### Pre-treatment cell cycle distributions and drug-induced cell cycle effects

As expected, we found that drug-induced changes in the G1, S and/or G2 fractions were dependent on the drug and cell line: a decrease in GR from 1 to 0 (net arrest) was most strongly associated with a reduction in S-phase cells (Spearman r = 0.57, P-value < 0.0001) and accumulation of cells in G1 or G2 (Fig. 6a). The CDK4/6 inhibitors ribociclib and abemaciclib shifted cell cycle distribution from S phase to G1 reflecting inhibition of the G1/S transition^41^, whereas BMS-265246, a drug primarily targeting CDK1/2^63^ induced G2 arrest (Fig. 6a, Supplementary Fig. 6a-b).

**Figure 6:**
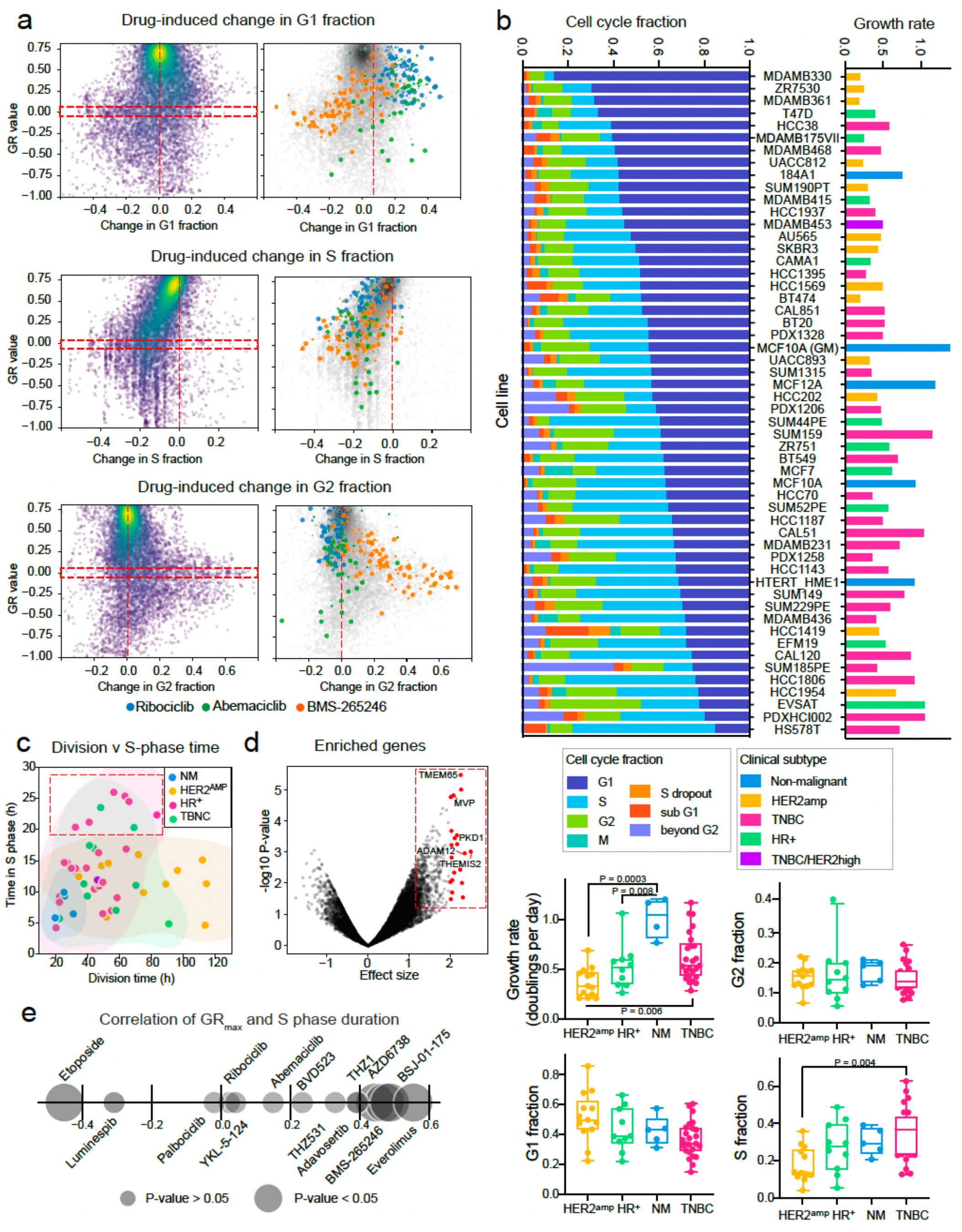
Cell cycle at baseline and in response to drug treatment. (a) Drug-induced change (compared to DMSO control) in the G1, S and G2 fractions with respect to GR value for all dose response data (left panels) and highlighted for ribociclib, abemaciclib and BMS-265246 (right panels). The size of the data point represents dose. (b) The baseline distribution throughout the cell cycle phases and growth rate in doublings per day for 58 breast cancer cell lines. Boxplots of the growth rate, and the fraction of cells at baseline in the G1, G2, and S phases of the cell cycle for the same cell lines separated by clinical subtype (n = 5 NM, 13 HER2^amp^, 13 HR^+^, 26 TNBC). The bottom and top of the box show the first and third quartiles, the bar within each box represents the median value and error bars represent the range of values. All cell lines were pulsed with EdU for 1 h to identify those in S-phase. The P-values indicated are adjusted P-values from 2-way ANOVA tests with Tukey’s correction for multiple comparisons. (c) Duration of S-phase with respect to division time for 58 breast cancer cell lines, colored by clinical subtype. The dashed red box indicates S-phase duration > 20 h. (d) Genes enriched in TNBC cells with S phases > 20 h relative to those with shorter S-phases. The 25 genes with the largest effect sizes are highlighted in red. (e) Spearman correlation between the duration of S-phase and the GRmax of the dose response curves across all cell lines for the drugs indicated.

The most remarkable feature of the data was not the drug response *per se* but the wide diversity of proliferation rates (reported here in divisions per day) and cell cycle distributions in the absence of drug exposure. Doubling times, as measured in DMSO-only cultures, varied ∼7-fold (from 17 h to 114 h per doubling) with HER2^amp^ lines the slowest growing (median 0.33 doublings/day) and NM cells the fastest growing (median 0.95 doublings/day) (Fig. 6b). This wide range of division times, a known confounder in comparative studies that use relative viability metrics like IC50 or AUC, highlights the importance of using GR metrics or similar methods to mitigate growth rate bias.^20, 21^ Across all lines, G1 fraction exhibited a significant negative correlation with division rate and S-phase exhibited a positive correlation (Spearman r = −0.58 and r = 0.56, respectively; P-values < 0.001; Supplementary Fig. 6c-d). However, these anticipated relationships masked dramatic variation in basal cell cycle state: under conditions of normal growth, G1 fraction varied from 15% to 86%, G2 fraction from 6% to 40%, and S phase fraction from 4% to 63% with no obvious association with subtype. The distribution of cell cycle states in actual tumors has also been shown to vary within and across cancers^37^ and clearly warrants additional study.

We do not yet understand, in molecular terms, why cell division varies so widely across breast cancer cell lines. To demonstrate the potential for combining Dye Drop results with existing expression data to study the cell cycle we looked at cells with extended S phase duration. Slow DNA replication and sensitivity to DNA damaging agents are features of cells exhibiting replication stress^64^. When we compared S-phase duration to division time for all cell lines, we found that a subset of HR^+^ and TNBC cells had S-phases longer than 20 h, as compared to a median of 11.5 h (Fig. 6c, red dashed box). Using Cancer Cell Line Encyclopedia (CCLE)gene expression data^65^ we found that the expression of genes such as PKD1, a marker of poor metastasis-free survival^66^ (Fig. 6d) and MVP, which is linked to chemoresistance (effect size > 2, t-test P-value < 0.05)^67^ was enriched in TNBC lines with extended S phases (as compared to all other TNBC lines). GSEA of the 25 most differentially expressed genes revealed upregulation of ‘*Wnt-activated receptor activity*’ (GO:0042813) and ‘*sphingosine-1-phosphate receptor activity*’ (GO:0038036) (**Supplementary Table 2**) in cell lines with extended S phases. Dysregulation of Wnt signaling is common in TNBC^68^ and both WNT and sphingosine-1-phosphate pathways are implicated in metastasis^55^. Moreover, when we examined the correlation between S-phase duration (in the absence of drug) and GR_max_, we observed significant correlation between time spent in replication and efficacy of five drugs: AZD6738, which inhibits ATR, a kinase that detects and responds to replication stress (Spearman r = 0.44, P-value = 0.03), adavosertib, which inhibits WEE1, the G2/M checkpoint kinase (Spearman r = 0.39, P-value = 0.05), and three drugs (BSJ-01-175 Spearman r = 0.49, P-value = 0.01; THZ1 Spearman r = 0.39, P-value = 0.01; and THZ531 Spearman r = 0.32, P-value = 0.1) that target CDK12/13, a known regulator of DNA damage repair and Wnt pathway genes^70^ (Fig. 6e, Supplementary Fig. 5e). These data suggest that the differences in baseline and drug-induced cell cycle states measured by Deep Dye Drop assays can serve as a starting point for identifying genes associated with a disordered cell cycle and responsiveness to specific drugs, such as those that target DNA damage pathways.

## DISCUSSION

In this paper we describe the development and testing of a family of Dye Drop methods for performing reliable and efficient high-throughput multi-well plate assays of viable and fixed single-cells. We show that gentle displacement of small volumes of liquid using a series of solutions having increasing concentrations of iodixanol, an inert chemical approved for use in humans, makes it possible to perform multi-step protocols without disturbing dying or weakly adherent cells. This improves the accuracy and reproducibility of simple LIVE/DEAD assays with the additional benefit of reducing costs by using smaller reagent volumes. However, the most valuable feature of Dye Drop is that it greatly facilitates multi-step live-cell measurements, such as EdU-incorporation, in 384 well plates while also enabling follow-on assays of fixed-cells using immunofluorescence. By measuring the responses of a panel of 58 breast cancer cell lines to 67 small molecule drugs (∼ 4,000 nine-point dose response curves), we demonstrate that these benefits of Dye Drop can be achieved at scale, even by individuals with only a few days of training on the method.

Dye Drop is flexible – different types of reagents (i.e., dyes, antibodies, etc.) can be combined in a single experiment and various antibodies can be used to quantify relevant metrics like DNA focus formation, MOMP, cell flattening etc. As a result, Dye Drop can capture morphological changes that cannot be readily detected using well-average methods or flow cytometry. Both manual multi-channel pipettes and automated reagent dispensers can be used in Dye Drop, and once cells are fixed, conventional plate washing is possible. This makes the method suitable for smaller research groups and also for core facilities. Dye Drop can also be coupled with multiplexed imaging (i.e., CyCIF^71^) so that 10-15 single-cell measurements can be performed on each well. In this case, it is possible to separate Dye Drop assays and CyCIF by several weeks, allowing the CyCIF antibody panel to be customized based on preliminary findings and to focus high-plex assays on the most informative conditions. We conclude that Dye Drop and its variants constitute a flexible and extendable set of methods for efficiently and accurately performing a wide variety of cell-based assays in 384 well plates.

Standardized data analysis pipelines are important for ensuring the accuracy and reproducibility of multi-parametric assays^16^. We have therefore developed a set of computational routines for designing and performing drug dose-response assays using the Dye Drop method (Supplementary Fig. 7, see online Methods). These are combined into a single tool box with scripts we previously developed for computing GR metrics^72^. Dye Drop dose-response analysis also features a series of “flags” that alert users when experimental design criteria and results such as the number of controls, the dose range, and the accuracy of curve fitting are suboptimal. Using these methods, Dye Drop can be used to study a range of acute drug-induced phenotypes including cell cycle arrest, MOMP, cell death, changes in cell morphology, induction of DNA damage foci etc., as well as time-dependent changes in GR values associated with adaptive drug resistance.

Simply counting viable cells or measuring a well-average surrogate such as ATP level (most commonly by assaying ATP levels in cell extracts using CellTiter-Glo®) does not distinguish cell cycle arrest (cytostasis) from cell killing (cytotoxicity). However, using a simple ODE-based model of cell cycle progression and Dye Drop data makes it possible to decompose the cytostatic and cytotoxic components of a drug response (GR^S^ and GR^T^) based on a single on-treatment timepoint (the collection of time-dependent response data enables more sophisticated decomposition). We have used this to study conditions where a drug causes GR ∼ 0 and there is no net cell growth. Under these conditions, we find that a state of balanced cell proliferation and death is about twice as common than true cytostasis (cell cycle arrest). Moreover, we indirectly infer that many drug responses are time-dependent, either involving adaptation or delayed cell killing (although proof that this is true will require additional time-dependent measurements). Finally, we find that the dose-dependence of GR^S^ and GR^T^ vary dramatically by cell line and drug. Some drugs — the FDA-approved CDK4/6 inhibitors, ribociclib and palbociclib for example — differ widely in potency (GEC^S^_50_) across a cell line panel but elicit little or no cell killing. In contrast, drugs such as the CDK1/2/5/9 inhibitor dinaciclib (whose clinical development ended in phase 2), have nearly the same potency in all cell lines and differ instead in cytotoxicity (GR^T^_max_). Drugs such as abemaciclib differ in both potency and efficacy^73^. These phenomena have not previously been described and their significance as yet unknown, but we note that abemaciclib is the most clinically active of the approved CDK4/6 inhibitors^74^, whereas dinaciclib and alvocidib^75^ are associated with serious toxicity in humans. We speculate that variation in both efficacy and potency, as exhibited by abemaciclib or the tool compound YKL-5-124, may be a property of an ideal cancer therapeutic.

The breast cancer lines used in this study vary up to 7-fold in their doubling times under normal growth conditions and the distributions of cells across the cell cycle also differ widely. Non-malignant cells are the fastest growing on average and HER2^amp^ cells the slowest growing (non-malignant cells also have the lowest rates of cell death in the absence of drug, median ∼2%). G1 and S-phases are the most variable across all lines, varying from 15% (HS578T cells) to 86% (MDAMB330) G1 fraction and 4% (MDAMB175VII) to 63% (HS578T) S phase fraction. The origins of this remarkable variability are not known but likely include the presence of recurrent mutations in genes that ensure the fidelity of DNA synthesis and repair (e.g. p53 and BRCA1/2) and the consequent replication stress^64^. Subtype-specific differences in proliferation demonstrate the importance of normalizing drug response to division rates using GR metrics or similar approaches. When such normalization is performed, multiple statistically significant genetic associations between drug response in cell lines and human patients can be identified; most of these are obscured by use of conventional IC50 metrics. Use of GR data also makes clear that non-malignant lines are not, on average, more resistant to approved anti-cancer drugs than cancer cell lines and that this criterion should not be used as a measure of tolerability in pre-clinical drug screens.

Even though imaging-based high content screening has been around for several decades, well average CellTiter-Glo® assays remain the most common measurement performed in cell-based drug discovery screens – particularly in oncology. The perceived simplicity and low cost of CellTiter-Glo® is the likely explanation for its popularity, but it yields only limited data that is not always interpretable as viable cell number. Our calculations show that reagent costs for Dye Drop and Deep Dye Drop assays are actually lower than those for CellTiter-Glo® assays (performed according to manufacturer’s instructions). When the additional costs of high-quality imaging plates and increased labor are factored in, Deep Dye Drop and CellTiter-Glo® are similar in cost. However, any of these assays represent only a small fraction (∼20%) of the total cost of performing a cell-based profiling or screening study at scale. Thus, Dye Drop methods represent a substantially more favorable balance between throughput, information content, and cost compared to current approaches. For chemistry campaigns or annotation of known bioactive collections (for which response rates are high), Deep Dye Drop plus CyCIF^71^ is likely the best approach; for large-scale, low hit-rate rate screens of small molecules, siRNA, or CRISPR-Cas9 libraries, minimal Dye Drop assays may be preferred (followed by re-screening hits with higher content assays). In both cases, Dye Drop assays provide new insight into response phenotypes and their relationship to doses that have been difficult to study at scale but are highly relevant to the use of small molecule drugs as research tools as well as their development into human therapeutics.

## METHODS

### Cell culture

Cell lines were maintained in their recommended growth medium and culture conditions as detailed in **Supplementary Table 3**. Conventional cell lines were identity verified by STR profiling, and newly established cell lines were STR profiled to ensure they were unique and to set benchmarks for future reference.

### Screening

Drugs were arrayed in nine-point half-log dilution series in 384 well library plates (**Supplementary Table 4**). The identity and purity of the drugs were verified by LC-MS. Each daughter plate contained 10 µl per well, and was thawed a maximum of 12 times. Cells were seeded in 384 well CellCarrier or CellCarrier ULTRA plates (Perkin Elmer, Waltham, MA) with a multidrop combi liquid dispenser at the densities listed in **Supplementary Table 3**, and allowed to adhere for 24-36 hours prior to drug treatment. Drugs were delivered from library plates via pin transfer with a custom E2C2515-UL Scara robot (Epson, Long Beach, CA) coupled to stainless steel pins (V&P Scientific, San Diego, CA) at the ICCB-Longwood Screening Facility, or from stock solutions with a D300 digital drug dispenser (Hewlett-Packard, Palo Alto, CA). At the time of drug delivery, replicate plates were stained and fixed to serve as time=0 reference data for GR-based calculations, and 72 hours later treated plates were stained and fixed according to the Dye Drop or Deep Dye Drop protocol (see below).

### Dye Drop assay

Cells, in 384 well plates, were stained by adding 15 µl of a staining solution per well: 1 µg/ml Hoechst 33342 (Thermo Fisher, Waltham, MA) and 1:2000 LIVE/DEAD far red dye (LDR) (Thermo Fisher, Waltham, MA) in 10% OptiPrep™ (Sigma, St. Louis, MO) in PBS (Corning, Glendale, AZ). After thirty minutes at room temperature (RT), cells were fixed by adding 20 µl of 4% formaldehyde (Sigma, St. Louis, MO) in 20% OptiPrep™ in PBS per well and incubating for 30 minutes. After fixation, the wells were aspirated and filled with 80 µl PBS, plates were sealed with foil adhesives (BioRad, Hercules, CA) and stored at 4°C until imaged. A 16-channel automatic multi-pipette was used for the addition of the stain and fix solutions, all aspirate steps and other dispense steps were performed with an EL406 washer equipped with a 96-channel head (Biotek, Winooski, VT).

### Deep Dye Drop assay

15 µl of a 10% OptiPrep™ solution in PBS containing 10 µM EdU (Lumiprobe, Waltham, MA) and 1:2000 LDR was added cell in each well of a 384 well plate and incubated for one hour at 37°C. Cells were then fixed in 20 µl/well 4% formaldehyde in 20% OptiPrep™for 30 minutes at RT. The duration of the EdU pulse can be adjusted depending on the division time of the cell line or on the experimental conditions, however, in our experience a one-hour pulse of EdU at 37°C is sufficient for most conventional cell lines. Following fixation, the wells were aspirated using an EL406 washer. 15 µl of cell permeabilization solution, 0.5% Triton-X100 (Sigma, St. Louis, MO) in 10% OptiPrep™, was then added per well at room temperature (RT) followed, 20 minutes later, by 20 µl of Click chemistry solution (2 mM copper sulfate, 4 µM Sulfo-Cy3 azide (Lumiprobe, Waltham, MA), 20 mg/ml ascorbic acid) in 20% OptiPrep™ to fluorescently label the incorporated EdU. After 30 minutes at RT, once the Click reaction was complete, the wells were aspirated, and 40 µl of Odyssey blocking buffer (LI-COR Biosciences, Lincoln, NE) was added per well for a minimum of one hour at RT, or up to overnight at 4°C to block the cells for immunofluorescence labeling. Next, an Odyssey blocking buffer solution containing 10% OptiPrep™ and 1:2000 Alexa 488-conjugated anti-phospho-histone H3 (S10) antibody (pH3) (clone D2C8, Cell Signaling Technology, Danvers, MA) to label M-phase cells and 2 µg/ml Hoechst 33342 to stain all nuclei, was dropped onto the cells and incubated overnight at 4°C. The overnight incubation ensures that the Hoechst staining saturates enabling accurate, image-based quantitation of DNA content. Post incubation, the plates were washed once with 0.01% PBS-Tween (Thermo Fisher, Waltham, MA) and twice with PBS, leaving a final volume of 80 µl of PBS in each well. The plates were then sealed with foil adhesives (BioRad, Hercules, CA) and kept at 4°C until imaged. As above, a 16-channel automatic multi-pipette was used for the addition of all stain and fix solutions, and all aspirate steps and other dispense steps were performed with an EL406 plate washer.

### Microscopy and feature extraction

Image acquisition was performed with either an Operetta (Perkin Elmer, Waltham, MA) or an ImageXpress Micro-Confocal (IXM-C) (Molecular Devices, San Jose, CA) high throughput microscope using a 10x objective. Six fields of view per well were acquired with the Operetta, and four with the IXM-C to achieve full well coverage. Both systems were equipped with robotics to enable continuous imaging 24 hours/day. Cell segmentation was performed with Columbus (Perkin Elmer, Waltham, MA) or MetaXpress (Molecular Devices, San Jose, CA) depending on the system used for image acquisition. In both cases, nuclei were segmented based on their Hoechst signal, a ring was drawn around each nucleus, and the average intensity of each stain was measured in each mask. For segmentation with Columbus, ‘Find Nuclei’ method ‘B’ was used to identify nuclei with the following settings: Common Threshold 0.5, Area > 50 µm^2^, Split Factor 7, Individual Threshold 0.4, and Contrast > 0.1. ‘Select Cell Region’ method ‘Resize Region [%]’ was used to draw a ring around each nucleus with the following settings: Region Type Nucleus Region, Outer Border −100%, Inner Border −30%. For segmentation with MetaXpress, nuclei were found using the ‘Find Round Objects’ module with minimum and maximum width thresholds of 8.2 µm and 23.4 µm, respectively, and an intensity above background threshold of 1366. The ‘Grow Objects Without Touching’ module was used to draw a 4 pixel ring around each nucleus. To account for local changes in background intensity, the ring intensity was subtracted from the nuclear intensity. The average nuclear Hoechst intensity was multiplied by the nuclear area to obtain the DNA content measurement.

### Validation of other antibodies and five channel Deep Dye Drop

Cells were seeded, treated and subjected to the Deep Dye Drop protocol as described above with the following modifications. Alternate antibodies were used in the place of pH3: actin (1:500, Cell Signaling Technologies, Danvers, MA), 53BP1 (1:500, BD Biosciences, Franklin Lakes, NJ), cytochrome C (1:200, Invitrogen, Carlsbad, CA), pRb (1:500, Cell Signaling Technologies, Danvers, MA). For five channel Deep Dye Drop, the protocol was followed as above, and cells were incubated overnight in the presence of pH3(S10) (1:2000, Cell Signaling Technologies, Danvers, MA) and 53BP1 (1:500, BD Biosciences, Franklin Lakes, NJ) primary antibodies. The cells were then washed and incubated with 1:2000 secondary donkey-anti-rabbit Alexa 488 and goat-anti-mouse Alexa 750 antibodies in Odyssey buffer for one hour at RT.

### Cyclic immunofluorescence

MCF7 and MCF 10A cells were seeded 15,000 and 8000 cells per well in two 96 well plates (Corning, Glendale, AZ) and allowed to adhere for 24 h prior to treatment with ribociclib, BMS-265246 or DMSO for 72 h. Cells were then stained and fixed according to the Deep Dye Drop protocol, the click-chemistry EdU labeling step was omitted on plate one. Cells were then imaged, plate two was washed once with EDTA (10 mM in PBS), incubated at RT for 2 h. in 10 mM EDTA (60 µl/well) and washed three times with PBS. Both plates were then bleached for 1 h (60 µl per well of 3% (wt/vol) H_2_O_2_, 20 mM NaOH in PBS) exposed to light, washed three times with PBS and subjected to three rounds of cyclic immunofluorescence.^35^ Plate one was then bleached again, washed three times with PBS, and the EdU was labeled per the Deep Dye Drop protocol. Hoechst (1 µg/ml) was included in all staining rounds. Image registration was performed with ASHLAR and nuclear segmentation and extraction of intensity features for each channel was performed with MCMICRO^76, 77^. Uniform Manifold Approximation and Projection (UMAP) was applied to the single cell intensity data for all antibody markers, EdU and the first instance of Hoechst staining (11-plex data, see **Supplementary Table 5**) using the umap library in python p).

### Data analysis

Analysis of the single cell level feature data was performed automatically with custom scripts (see Data and code availability below). The detailed computational protocol for gating measured signal intensities is available at https://github.com/datarail/DrugResponse/wiki. Briefly, The LDR, EdU, and pH3 intensities were log transformed and smoothed using a kernel density estimate (KDE) function. A peak finding algorithm was used to identify the global minima of the KDEs. The minima were used to set thresholds above which the cells were classified as dead (based on LDR), in S-phase (based on EdU) or in M-phase based on pH3 intensity. The integrated Hoechst intensity was used to quantify DNA content to discriminate between cells in the G1 and G2 phases of the cell cycle. Cells that were negative for EdU but had intermediate DNA content between thresholds set for G1 and G2 were classified as ‘S dropout’. Cells positive for LDR but that no longer harbored any Hoechst signal were scored as ‘corpses’ and were included in the total dead cell count. Spatial analysis was performed by using X-Y pixel coordinates to calculate the shortest Euclidean distance between every S-phase cell to its nearest S-phase cell and also the shortest distance between each S-phase cells and the nearest cell in any other phase. The distribution of the distances between the two groups were compared. Boxenplots to visualize the data were generated using https://seaborn.pydata.org/generated/seaborn.boxenplot.html.

### Standard GR, GR static, and GR toxic

Standard GR value are calculated as defined previously based on the number of viable cells (Hoechst positive and LDR negative).^17^ Quantification of the cytostatic and cytotoxic components of the response relies on a simple model of population growth with live cells, *x*, growing exponentially at a doubling rate *k_s_* while dead cells, *d*, are dying proportionally to *x* at a rate [*log(2) k_d_*]. Consequently, the population model is ruled by the ODE system:

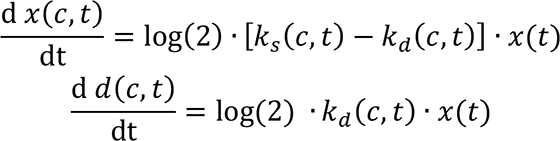

Note that the term log(2) is a factor to convert doubling rate into growth rate and allows us to solve this ODE system for *k_s_* and *k_d_* as:

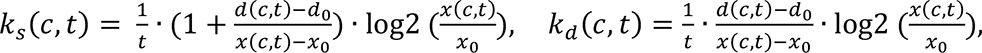

Where:

- *x(c,t)* is the number of viable cells at time t at drug concentration c
- *d(c,t)* is the number of dead cells at time t at drug concentration c
- *x_0_ = x(0,0)*, the number of live cells at the beginning of the treatment
- *d_0_ = d(0,0)*, the number of dead cells at the beginning of the treatment
- *t* is the continuous time in an experiment over which responses are integrated

The rates *k_s_(c,t)* and *k_d_(c,t)* can be normalized by the untreated control rates and mapped to range of [0, 1] and, respectively, [-1, 0] defining *GR_s_*(*c*) and *GR*_T_(*c*) as:

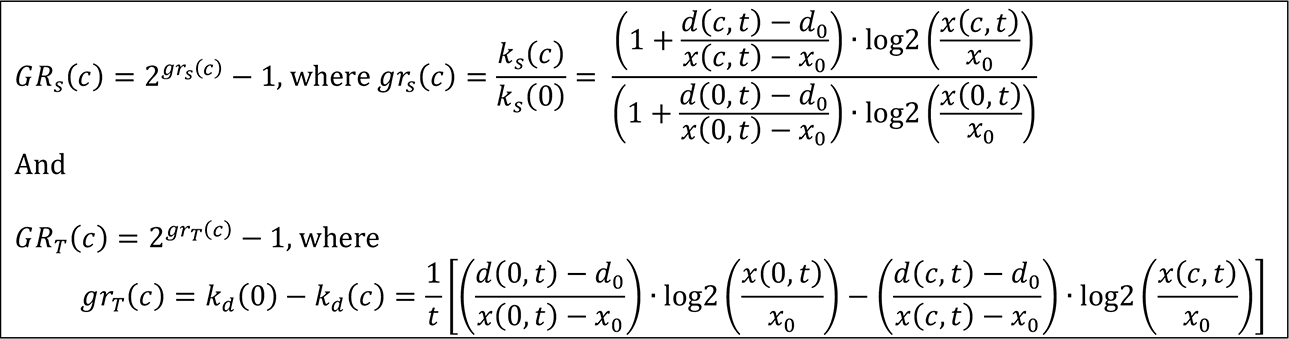

Note that we set *d*(*c*, *t*) – *d*_0._ to 1 if below 1 because dead cells are meant to be cumulative over the course of the experiment and that a numerical approximation based on a Taylor expansion is used if *x*(*c*, *t*) ≅ *x*. (see code below).

GR values are related to *GR_S_* and *GR_T_* values as follows:

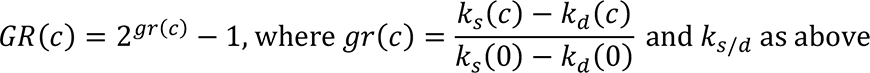

With the simplification of *k_d_* (0) = 0:

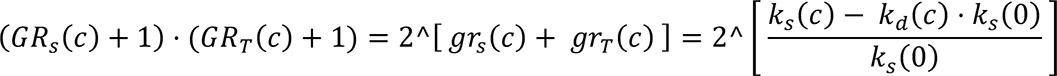

Thus we can see that:

(*GR_S_*(*c*) + 1) ⋅ (*GR_T_*(*c*) + 1) ≈ (*GR*(*c*) + 1) in cases where *k_d_*(*c*) ≫ 0 or *k_s_*(0) ≅ 1.

### Gene set enrichment analysis in cell lines with extended S-phase duration

Gene expression data for TNBC cell lines in our study were downloaded from the Broad Cancer Cell Line Encyclopedia 21Q2 data release (https://depmap.org/portal/download/)^78^. Cell lines were separated into those with S phase longer than 20 h and all others. Cohen’s d was used to measure the effect size for all genes. The 25 genes with the largest effect sizes were entered in Enrichr (https://maayanlab.cloud/Enrichr/)^79,80^ to identify enriched Gene Ontology (GO) Molecular Function terms.

### Data availability

All data and code as well as additional relevant resources are available at https://labsyspharm.github.io/dye-drop-microsite/. This site will continue to be updated will new results, tools, and links to complementary projects in our lab. GR, cell death, and cell cycle results for the breast cancer profiling dataset presented in this paper are available for download from synapse (https://www.synapse.org/#!Synapse:syn26133007) and can be browsed online: https://labsyspharm.shinyapps.io/HMSLINCS_BRCA_Browser/. Additional data corresponding to the figures presented in this paper are also available under the same synapse ID. 21Q2 public CCLE gene expression data were used for Fig. 6d and were obtained from: https://depmap.org/portal/download/. Detailed Dye Drop and Deep Dye Drop protocols are available on protocols.io: https://www.protocols.io/view/deep-dye-drop-protocol-96zh9f6.

### Code availability

All scripts needed to analyze single cell intensity data are available on github: github.com/datarail/DrugResponse/tree/master/python/cell_cycle_gating and a user guide detailing installation of these tools and their execution is available online: https://ddd-gating.readthedocs.io/en/latest/index.html. The code needed for GR calculations is on github: github.com/datarail/gr_metrics. A web resource for calculating GR values and metrics is also available at grcalculator.org where additional information is available. The modularity of the suite of tools means that each component can be used independently of the others, or jointly depending on the experiment and equipment available. Refer to Supplementary Fig. 7 for an overview of these resources and how the modules fit together.

## ACKNOWLEDGEMENTS

We thank ICCB-L for assistance with compound management and drug treatments. We thank M. P. Rout for the suggestion that OptiPrep™ be used for Dye Drop, N. Gray, for providing compounds, J. Tefft for editing the manuscript and J. Tefft and H. Xu for website development. This work was funded by NIH grants U54-CA225088, U54-HL127365 and U24-DK116204 and a generous gift from the Termeer Foundation.

## OUTSIDE INTERESTS

PKS is a co-founder and member of the BOD of Glencoe Software, a member of the BOD for Applied Biomath, and a member of the SAB for RareCyte, NanoString and Montai Health; he holds equity in Glencoe, Applied Biomath and RareCyte. PKS is a consultant for Merck and the Sorger lab has received research funding from Novartis and Merck in the past five years. Sorger declares that none of these relationships have influenced the content of this manuscript. KS is currently an employee of Bristol Myers Squibb, MH and LG are currently employees of Genentech, and MN is currently an employee of Ribon Therapeutics. The other authors declare no outside interests.

---

**Supplementary Figure 1:**
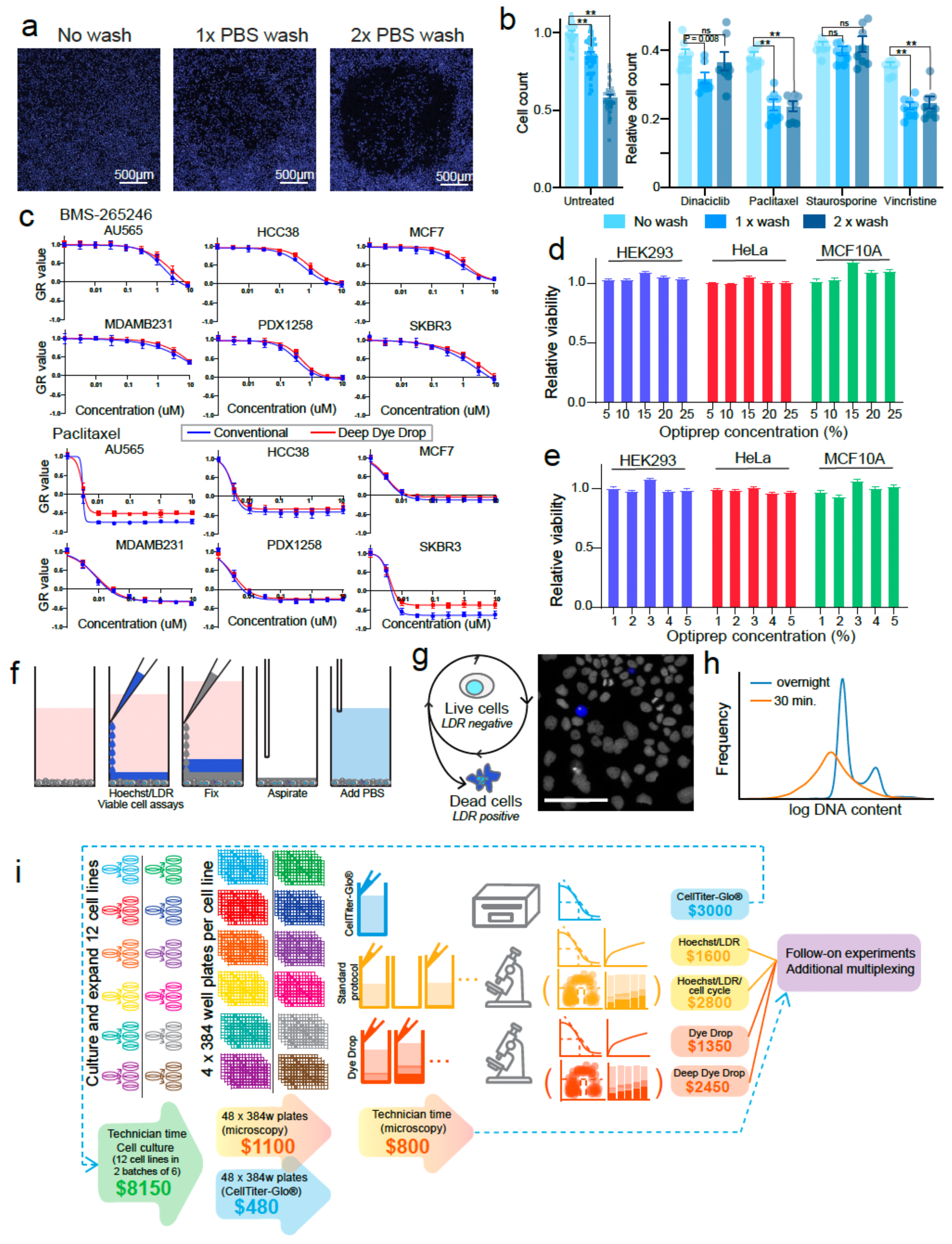
Validation of assay conditions and assay schematics. (a) Images of MCF 10A-H2B-mCherry cells in a well of a 384 well plate stained with Hoechst in OptiPrep™ without prior washing and following one or two PBS wash cycles with a robotic plate washer. (b) Consequences of one or two wash cycles prior to fixation on MCF 10A-H2B-mCherry cells untreated and treated with 0.1 µM dinaciclib, paclitaxel, staurosporine, and vincristine for 24 h. Error bars represent the standard error of the mean of eight technical replicates from one representative biological replicate, P-values shown are P-values from 2-way ANOVA tests corrected for multiple comparisons with Tukey’s method, ** indicates P-value < 0.0001, ns indicates not significantly different. (c) GR dose response curves for six breast cancer cell lines treated with BMS-265246 or paclitaxel for 72 h at the doses indicated and assayed by Deep Dye Drop (red lines) or by a conventional washing and staining protocol (blue lines). (d) Effects of a one hour pulse of increasing concentrations of OptiPrep™ on the viability of HEK293, HeLa and MCF 10A cells 24 h later. Error bars represent standard error of the mean of 200 technical replicates. (e) Effects of a 24 h exposure to increasing concentrations of OptiPrep™ on the viability of HEK293, HeLa and MCF 10A cells. Error bars represent standard error of the mean of 200 technical replicates. (f) Dye Drop protocol steps: Hoechst and LIVE/DEAD (LDR) dye are added in 10% OptiPrep™ followed by 4% formaldehyde in 20% OptiPrep™, the contents of the well are aspirated, and replaced with PBS. One well of a multi-well plate is depicted. (g) Schematic of Dye Drop staining and image showing cells stained with the Dye Drop protocol. Hoechst staining is gray-scale and LDR staining is blue. Scale bar is 100 µm. (h) DNA content quantified from untreated BT20 cells stained with the Dye Drop (Hoechst 30 min) and Deep Dye Drop (Hoechst overnight (o/n)) protocols. (i) Schematic diagram of the costs associated with running a 12 cell line, 30 drug profiling experiment by CellTiter-Glo®, Dye Drop, Deep Dye Drop or equivalent assays with a conventional protocol.

**Supplementary Figure 2:**
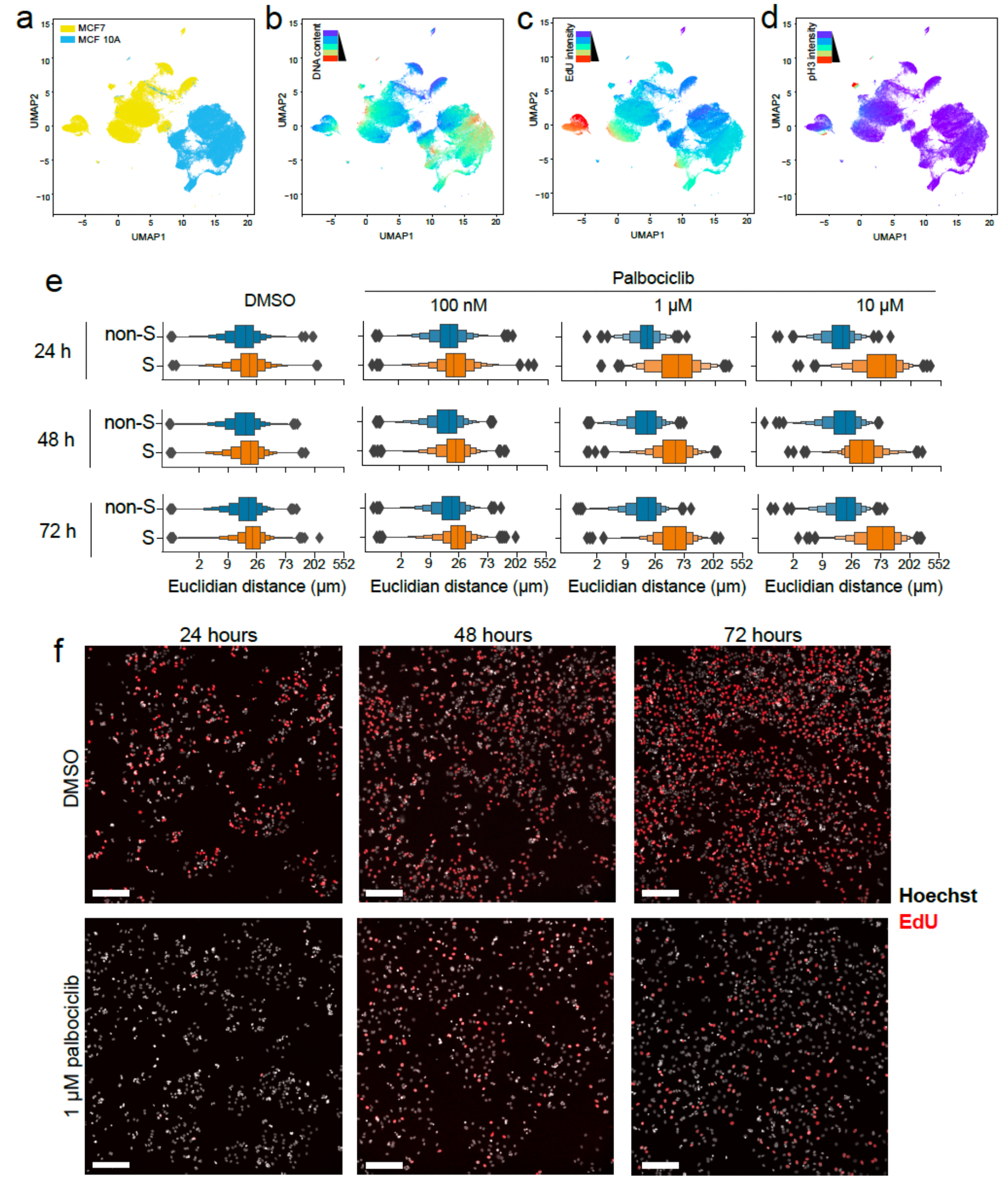
Extension of Deep Dye Drop assays. UMAP representation of MCF7 and MCF 10A cells treated with BMS-265246 (1 µM, 10 µM), ribociclib (10 µM) or DMSO stained with Deep Dye Drop and cyclic immunofluorescence colored by (a) cell line, (b) DNA content, (c) EdU intensity, and (d) phospho-histone H3 intensity. (e) Boxenplots showing the natural log Euclidian distance from S-phase cell to the nearest S-phase cell and between S-phase and the nearest cell assigned to any other cell cycle phase in a population of MCF7 cells treated with DMSO or palbociclib at 1 µM or 10 µM for 24, 48 and 72 h. The centerline in each plot is the median, each successive level outward contains half of the remaining data. (f) Representative images of MCF7 cells treated with DMSO or 1 µM palbociclib for 24, 48, or 72 h. Scale bars are 200 µm.

**Supplementary Figure 3:**
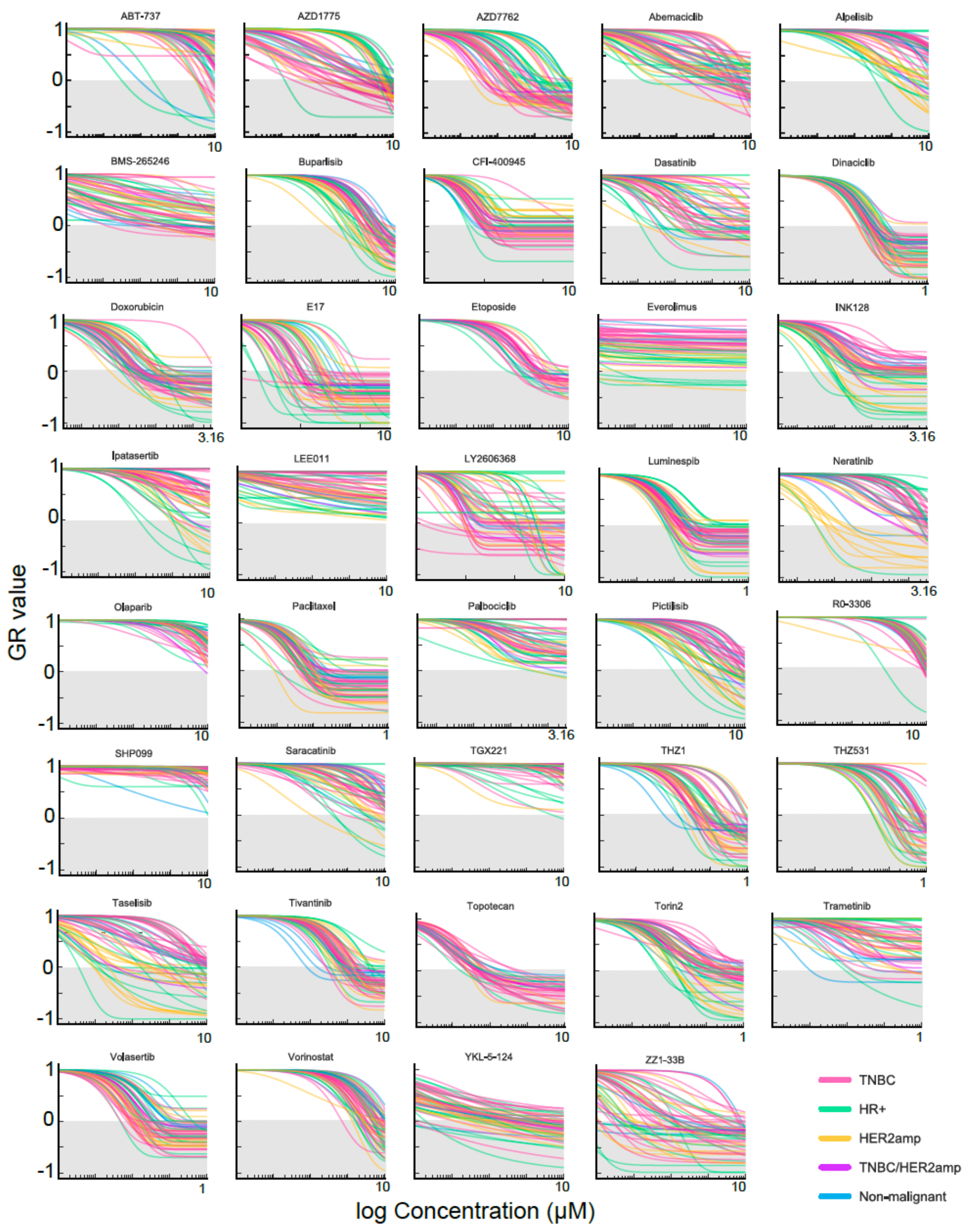
GR dose response curves for select drugs in 58 cell lines. GR-based dose response curves for 58 breast cancer cell lines treated with increasing concentrations of the drugs indicated for 72 hours. Cells were either stained with the Dye Drop or Deep Dye Drop assays. Each curve represents the fit to the average of three or four technical replicates, error bars are not shown for visual simplicity. Curves are colored by clinical subtype.

**Supplementary Figure 4:**
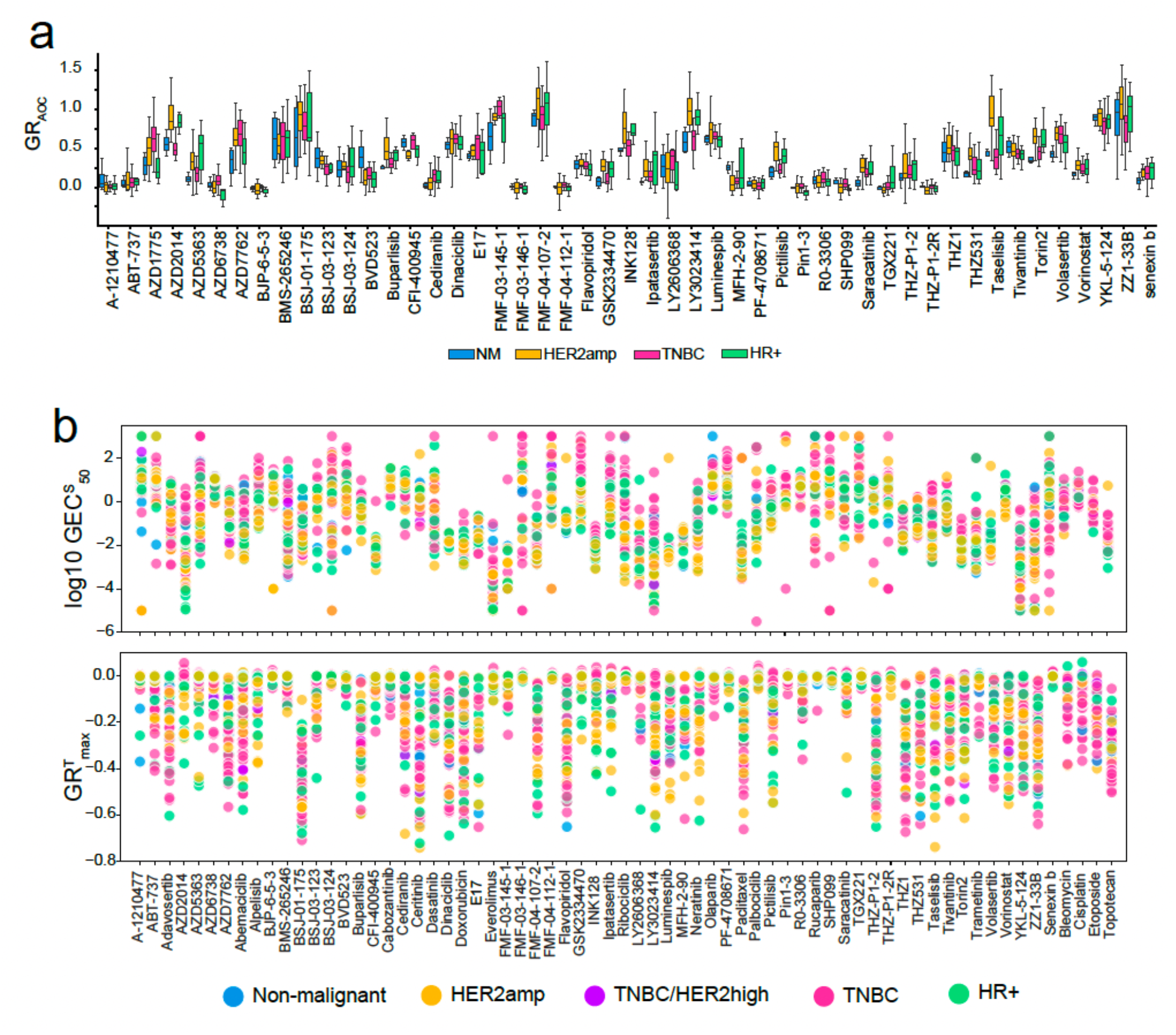
GR metrics definitions and results for 67 drugs in 58 breast cancer cell lines. (a) Area over the GR curves for the responses of non-FDA-approved drugs screened in 58 cell lines colored by clinical subtype (n = 5 NM, 13 HER2^amp^, 13 HR^+^, 26 TNBC). The bottom and top of the box show the first and third quartiles, the bar within each box represents the median value and error bars represent the range of values. (b) GEC^S^_50_ and (e) GR^T^_max_ metrics for 67 drugs in 58 breast cancer cell lines.

**Supplementary Figure 5:**
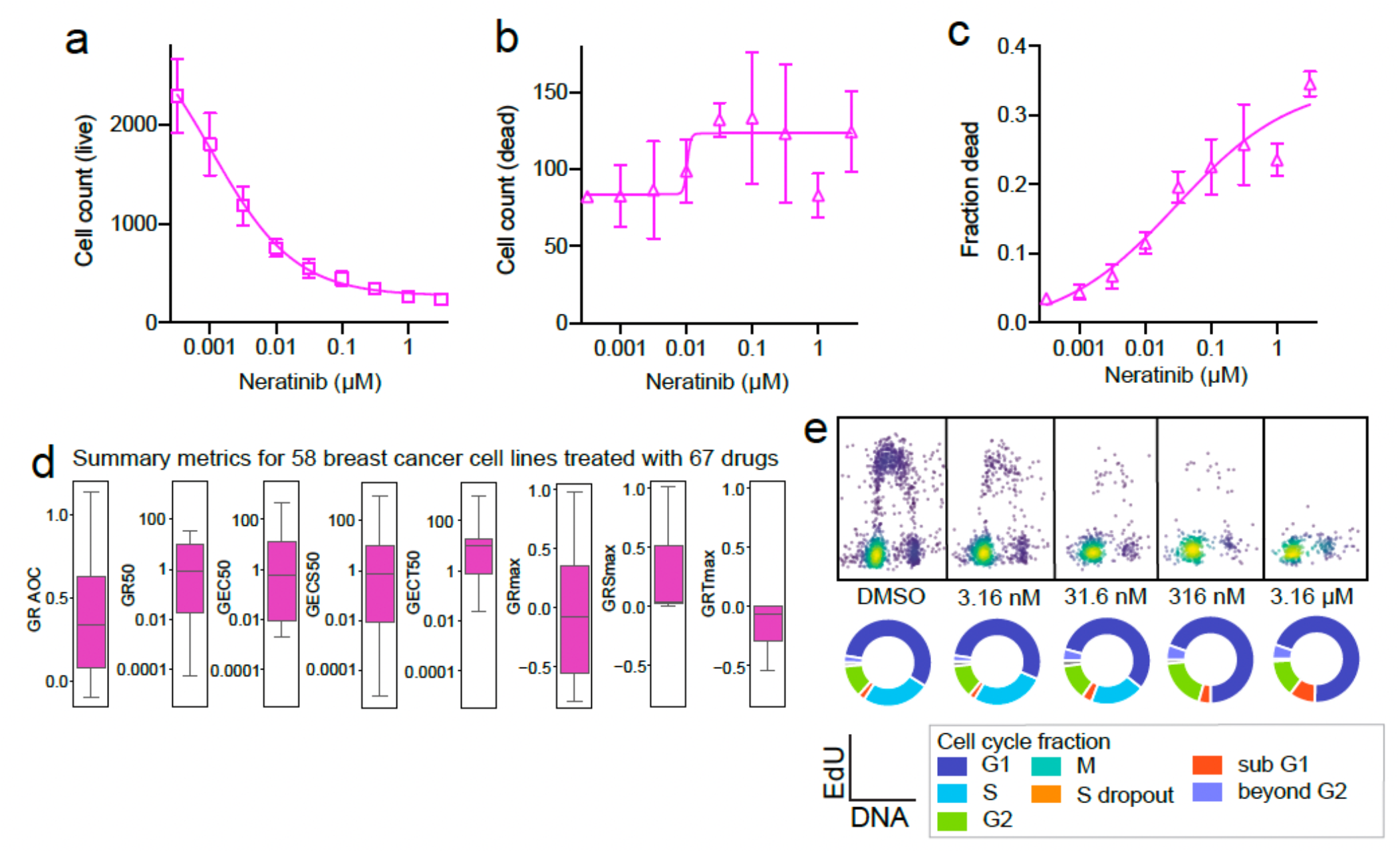
Neratinib response in AU-565 cells. (a) Live cell counts, (b) dead cell counts and (c) fraction of cells that are dead in AU-565 cells treated with neratinib at increasing concentrations for 72 h. Error bars represent standard deviation of the mean of technical quadruplicates. (d) GR metrics across 58 breast cancer cell lines treated with neratinib. The bottom and top of the box show the first and third quartiles, the bar within each box represents the median value and error bars represent the range of values. (e) Single cell EdU vs DNA content plots for the same conditions at the concentrations indicated with corresponding circular cell cycle fraction charts. All cells from a single well of a 384 well plate are shown.

**Supplementary Figure 6:**
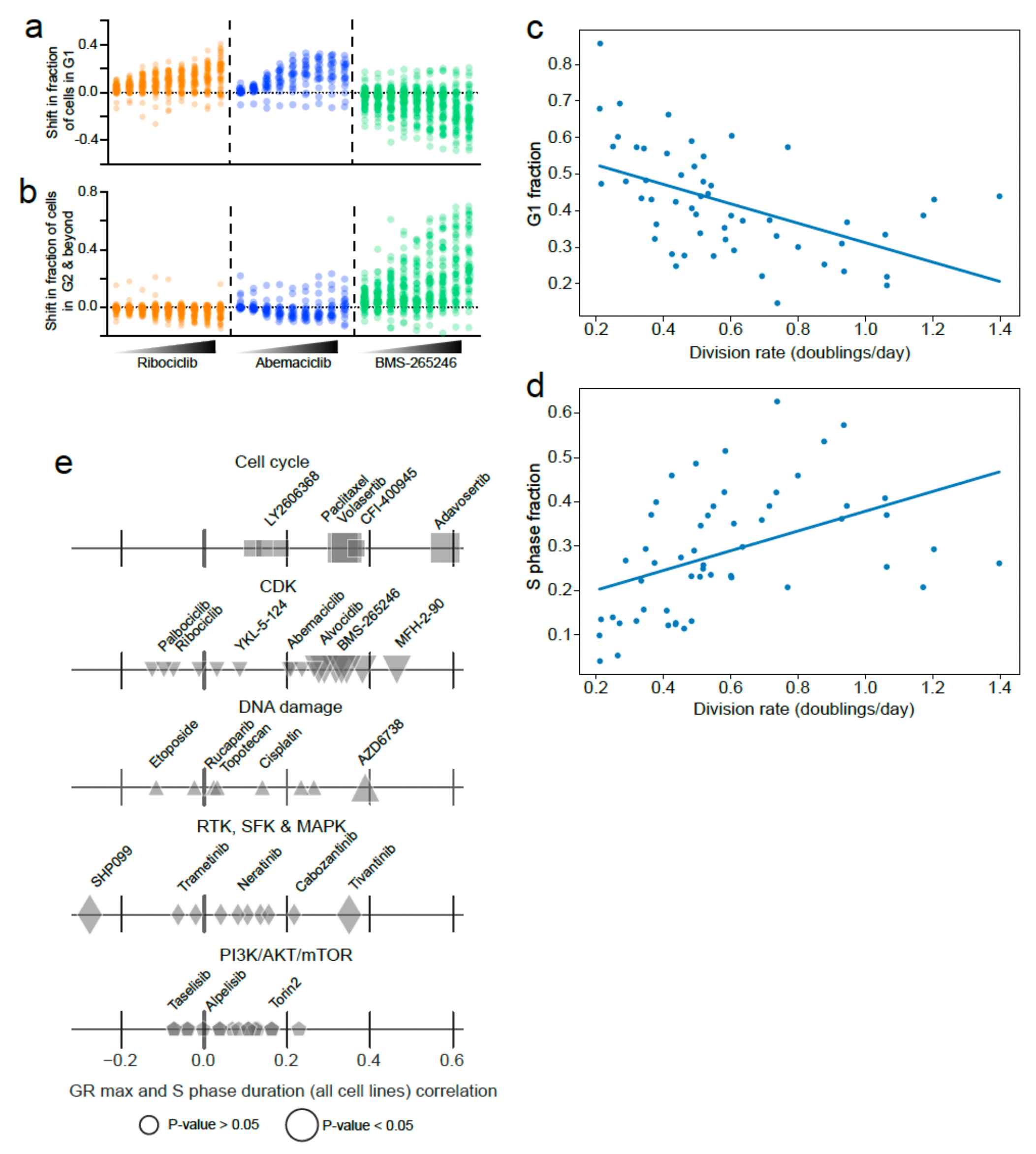
Relationship between baseline cell cycle distribution and drug response. (a) The effects of increasing concentrations of ribociclib, abemaciclib and BMS-265246 on the fraction of cells in G1, and (b) in G2 or with DNA content in excess of G2. (c) G1 and (d) S phase fraction with respect to division rate for 58 breast cancer cell lines. Lines of best fit are shown. (e) Spearman correlation between the duration of S-phase and the GRmax of the dose response curves across all cell lines for all drugs tested organized by pathway targeted.

**Supplementary Figure 7:**
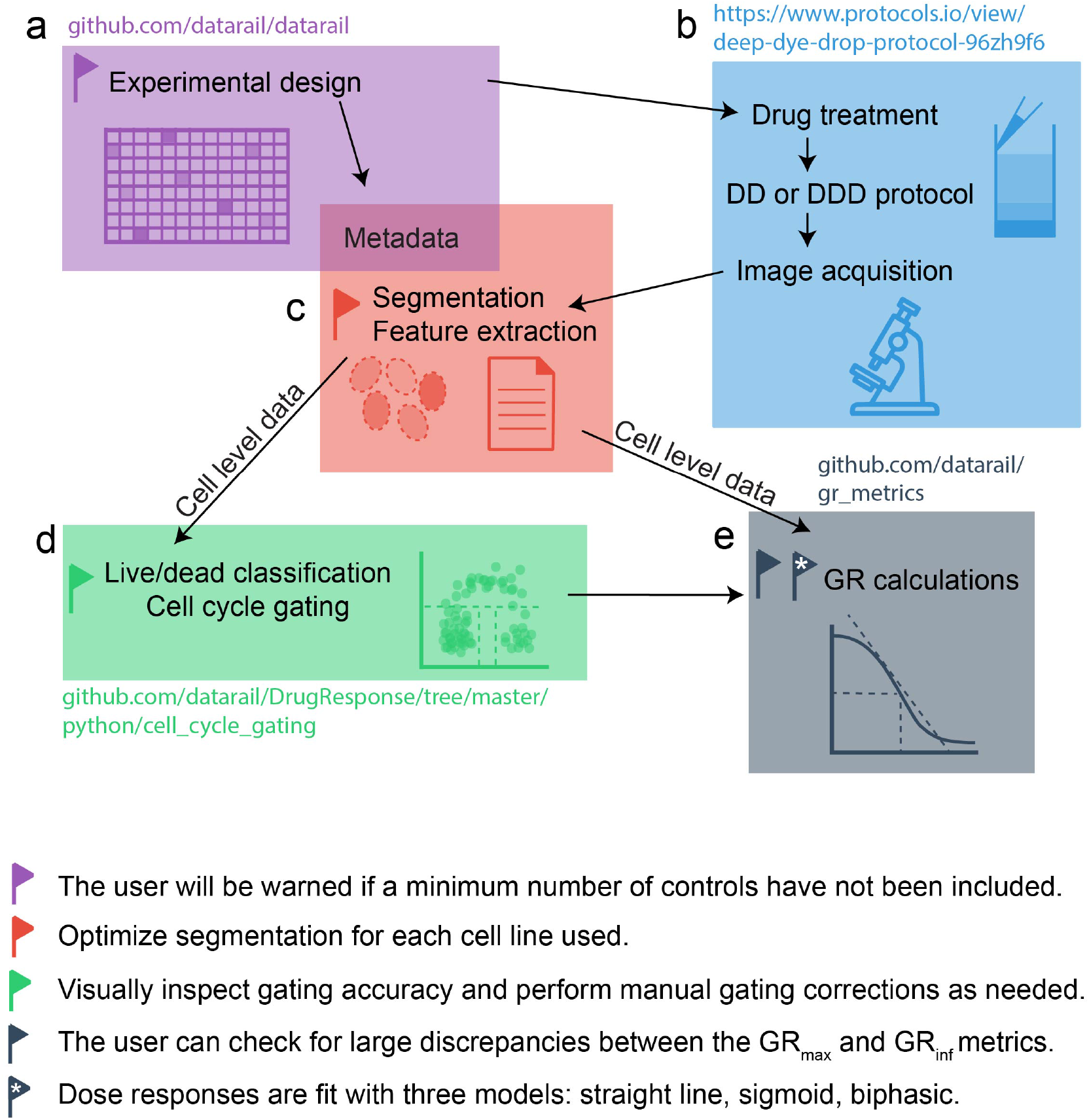
Overview of the components of a dose response experiment and available tools. (a) Automated experimental design Jupyter notebook for the randomization of dose response studies in multi-well plates. User input: experimental parameters (drugs, concentrations, time points, cell lines etc.); output: treatment file for a D300 digital drug dispenser and the associated metadata. The user is alerted if their experimental design does not include sufficient control wells (set at 8 per cell line). (b) Following drug treatment, cells are stained and fixed using the Dye Drop (DD) or Deep Dye Drop (DDD) assay, and images are acquired on any high throughput microscope. (c) Segmentation and feature extraction are performed and merged with the metadata. (d) Single cell level data can be gated automatically into the phases of the cell cycle. The user is presented with the gates overlaid on EdU versus DNA content scatter plots for visual inspection; should the gating be inaccurate it can be manually adjusted. Users have the option of having gates defined on negative control wells applied to treated wells, or of gating each well independently. (e) Well level data, either from feature extraction software, or summarized from single cell gating, are used to calculate GR values and fit GR metrics either using the online calculator (grcalculator.org), or a Jupyter notebook. (f) Stacked bar graphs for cell cycle, dose response curves (GR values, GR static, GR toxic and fraction dead), and summary metrics (per drug -cell line pair for each timepoint) are output. The suite of tools is modular, each component can be used independently of the others, or jointly depending on the experiment and equipment available.

