## Supplementary Tables for "Multiplexed and reproducible high content screening of live and fixed cells using the Dye Drop method"

**SUPPLEMENTARY INFORMATION**

**Supplementary Table 1:** Estimated costs associated with common dose response assays performed by an experienced technician in Boston, MA in 2022.

| **Cost per plate (summary)** | | | |
|---|---|---|---|
| **Assay** | **Reagents** | **Reagents + plates** | **Reagents + plates + labor** |
| **CTG** | $51.58 | $61.52 | $231.31 |
| **Hoechst/LDR** | $9.91 | $33.05 | $219.51 |
| **DD** | $5.19 | $28.32 | $214.78 |
| **Hoechst/LDR/EdU/IF** | $34.49 | $57.63 | $244.09 |
| **DDD** | $27.94 | $51.07 | $237.53 |

| **Cost for a 12 cell line, 30 drug screen (48 plates) (summary)** | | | |
|---|---|---|---|
| **Assay** | **Reagents** | **Reagents + plates** | **Reagents + plates + labor** |
| **CTG** | $2,475.60 | $2,952.72 | $11,102.72 |
| **Hoechst/LDR** | $475.84 | $1,586.44 | $10,536.44 |
| **DD** | $248.94 | $1,359.54 | $10,309.54 |
| **Hoechst/LDR/EdU/IF** | $1,655.50 | $2,766.10 | $11,716.10 |
| **DDD** | $1,340.95 | $2,451.55 | $11,401.55 |

| **Cost for a 12 cell line, 30 drug screen (48 plates) (detailed breakdown)** | | | | |
|---|---|---|---|---|
| **Personnel** | **Time** | **Rate** | **Notes** | **Total** |
| Technician 50% time 2 weeks (cell culture) | 40 h | $50/h | x 2 6 cell line batches | $4,000 |
| Technician 100% time 3 days plating, treating, staining/fixing | 24 h | $50/h | x 2 6 cell line batches | $2,400 |
| Technician time for microscopy-based assays (1 extra day) | 8 h | $50/h | x 2 6 cell line batches | $800 |
| **Consumables** | **Amount** | **Stock cost** | | **Total cost** |
| Growth media | 500 ml/cell line | $25/bottle | | $300 |
| Serum | 50 ml/cell line | $900/500 ml | | $1,000 |
| Plasticware/consumables | 1 sleeve of 10 cm plates/cell line | $350/case | | $350 |
| Miscellaneous | tips, pipettes | | | $100 |
| **CTG** | | | | **$8,150** |

| Microscopy assays | | | | | $8,950 |
|---|---|---|---|---|---|
| **Plates** | **Amount** | **Stock cost** | **Notes** | | **Total cost** |
| 384w cell culture plates | 48 total | $994/100 | CTG | | **$477** |
| 384w imaging plates | 48 total | $3702/160 | Microscopy | | **$1,111** |
| **Reagents** | **Components** | **Stock cost** | **Notes** | **Volume needed** | **Total cost** |
| **CTG** | | | | | |
| CellTiter-Glo (30 µl working volume) | CellTiter-Glo | $2063/500 ml | 1:1 | 600 ml | **$2,476** |
| **Conventional LIVE/DEAD assay** | | | | | |
| Stain (70 µl working volume) | Hoechst | $123/10 ml | 1:10000 | 130 µl | $1.60 |
| | LDR | $356/500 µl | 1:2000 | 650 µl | $463 |
| Fix | Formaldehyde | $176/L | 4% | 65 ml | $11 |
| | | | | | **$475.84** |
| **Dye Drop** | | | | | |
| Stain | Hoechst | $123/10 ml | 1:10000 | 30 µl | $0.37 |
| | LDR | $356/500 µl | 1:2000 | 150 µl | $107 |
| | Optiprep | $305/250 ml | 10% | 30 ml | $37 |
| Fix | Formaldehyde | $176/L | 4% | 43 ml | $8 |
| | Optiprep | $305/250 ml | 20% | 80 ml | $98 |
| | | | | | **$248.94** |
| **Conventional LIVE/DEAD + EdU + IF** | | | | | |
| Stain 1 | EdU | $30/10 ml | 10 µM | 1.3 ml | $3.90 |
| | LDR | $356/500 µl | 1:2000 | 650 µl | $463 |
| Fix | Formaldehyde | $176/L | 4% | 65 ml | $11 |
| Permeabilize | Triton X100 | $0.87/100 ml | 0.10% | 6 ml | $0.05 |
| Stain 2 | cy-3 azide | $140/ml | 4 µM | 600 µl | $84.00 |
| | CuSO4 | $1/5 ml | 2 mM | 6 ml | $1.20 |
| | Ascorbic acid | $50/100 g | 20 mg/ml | 12 g | $6.00 |
| Block | Odyssey | $116/500 ml | | 760 ml | $176.32 |
| Stain 2 | pH3 antibody | $408/100 µl | 1:2000 | 200 µl | $816.00 |
| | Hoechst 33342 | $123/10 ml | 1:5000 | 80 µl | $0.98 |
| | Odyssey | $116/500 ml | | 300 ml | $92.80 |
| | | | | | **$1,655.50** |
| **Deep Dye Drop** | | | | | |

| | | | | | |
|---|---|---|---|---|---|
| Stain 1 | EdU | $30/10 ml | 10 µM | 300 µl | $0.90 |
| | LDR | $356/500 µl | 1:2000 | 150 µl | $107 |
| | Optiprep | $305/250 ml | 10% | 30 ml | $37 |
| Fix | Formaldehyde | $176/L | 4% | 43 ml | $8 |
| | Optiprep | $305/250 ml | 20% | 80 ml | $98 |
| Permeabilize | Triton X100 | $0.87/100 ml | 0.10% | 3 ml | $0.03 |
| | Optiprep | $305/250 ml | 10% | 30 ml | $37 |
| Click labeling reaction | cy-3 azide | $140/ml | 4 µM | 400 µl | $56.00 |
| | CuSO4 | $1/5 ml | 2 mM | 4 ml | $0.80 |
| | Ascorbic acid | $50/100 g | 20 mg/ml | 8 g | $4.00 |
| | Optiprep | $305/250 ml | 20% | 80 ml | $98 |
| Block | Odyssey | $116/500 ml | | 760 ml | $176.32 |
| Immunofluorescence | pH3 antibody | $408/100 µl | 1:2000 | 150 µl | $612.00 |
| | Hoechst 33342 | $123/10 ml | 1:5000 | 60 µl | $0.74 |
| | Odyssey | $116/500 ml | | 300 ml | $69.60 |
| | Optiprep | $305/250 ml | 10% | 30 ml | $37 |
| | | | | | **$1,340.95** |

**Supplementary Table 2:** Gene Ontology Molecular Function terms enriched in in triple negative breast cancer cell lines with extended (>20 h) S-phases. Analysis performed with Enrichr (https://maayanlab.cloud/Enrichr/). P-value is from the Fisher exact test, and the adjusted P-value is corrected for multiple hypothesis testing with the Benjamin-Hochberg method.

| ***Term*** | ***P-value*** | ***Adjusted P-value*** | ***Odds Ratio*** | ***Combined Score*** |
|---|---|---|---|---|
| Wnt-activated receptor activity (GO:0042813) | 0.0002 | 0.007 | 133.5 | 1170.5 |
| phosphotransferase activity, alcohol group as acceptor (GO:0016773) | 0.004 | 0.044 | 23.7 | 131.3 |
| sphingosine-1-phosphate receptor activity (GO:0038036) | 0.006 | 0.044 | 208.0 | 1056.3 |
| syndecan binding (GO:0045545) | 0.006 | 0.044 | 208.0 | 1056.3 |
| lipid kinase activity (GO:0001727) | 0.006 | 0.044 | 208.0 | 1056.3 |
| serine-type endopeptidase activity (GO:0004252) | 0.008 | 0.044 | 16.8 | 81.9 |
| kinase activity (GO:0016301) | 0.009 | 0.044 | 15.7 | 74.7 |
| protein phosphatase binding (GO:0019903) | 0.010 | 0.044 | 14.3 | 65.3 |
| serine-type peptidase activity (GO:0008236) | 0.011 | 0.044 | 14.0 | 63.8 |
| intramolecular transferase activity, phosphotransferases (GO:0016868) | 0.011 | 0.044 | 104.0 | 467.2 |

| | | | | |
|---|---|---|---|---|
| bioactive lipid receptor activity (GO:0045125) | 0.011 | 0.044 | 104.0 | 467.2 |
| diacylglycerol kinase activity (GO:0004143) | 0.014 | 0.049 | 83.2 | 357.1 |
| kinase binding (GO:0019900) | 0.019 | 0.064 | 5.8 | 23.0 |
| antiporter activity (GO:0015297) | 0.025 | 0.076 | 43.8 | 161.9 |
| protein phosphatase 2A binding (GO:0051721) | 0.030 | 0.085 | 36.1 | 127.2 |
| acetyltransferase activity (GO:0016407) | 0.039 | 0.099 | 26.8 | 86.8 |
| chemoattractant activity (GO:0042056) | 0.039 | 0.099 | 26.8 | 86.8 |
| endopeptidase activity (GO:0004175) | 0.058 | 0.136 | 5.5 | 15.5 |
| cholesterol binding (GO:0015485) | 0.061 | 0.136 | 16.9 | 47.5 |
| calcium ion transmembrane transporter activity (GO:0015085) | 0.067 | 0.136 | 15.4 | 41.6 |
| secondary active transmembrane transporter activity (GO:0015291) | 0.068 | 0.136 | 15.1 | 40.6 |
| sterol binding (GO:0032934) | 0.072 | 0.136 | 14.1 | 36.9 |
| SH3 domain binding (GO:0017124) | 0.075 | 0.136 | 13.6 | 35.3 |
| PDZ domain binding (GO:0030165) | 0.076 | 0.136 | 13.4 | 34.5 |
| acyltransferase activity, transferring groups other than amino-acyl groups (GO:0016747) | 0.091 | 0.156 | 11.1 | 26.5 |
| calcium channel activity (GO:0005262) | 0.100 | 0.165 | 10.0 | 23.0 |
| cation channel activity (GO:0005261) | 0.116 | 0.184 | 8.5 | 18.4 |
| protein kinase binding (GO:0019901) | 0.131 | 0.189 | 3.4 | 6.8 |
| phosphatase binding (GO:0019902) | 0.133 | 0.189 | 7.3 | 14.8 |
| metal ion binding (GO:0046872) | 0.136 | 0.189 | 3.3 | 6.6 |
| protease binding (GO:0002020) | 0.138 | 0.189 | 7.1 | 14.0 |
| metallopeptidase activity (GO:0008237) | 0.141 | 0.189 | 6.9 | 13.5 |
| magnesium ion binding (GO:0000287) | 0.167 | 0.218 | 5.7 | 10.2 |
| protein heterodimerization activity (GO:0046982) | 0.210 | 0.266 | 4.4 | 6.9 |
| ubiquitin protein ligase binding (GO:0031625) | 0.284 | 0.347 | 3.1 | 3.9 |
| ATP binding (GO:0005524) | 0.295 | 0.347 | 3.0 | 3.6 |
| ubiquitin-like protein ligase binding (GO:0044389) | 0.299 | 0.347 | 2.9 | 3.5 |
| adenyl ribonucleotide binding (GO:0032559) | 0.320 | 0.354 | 2.7 | 3.1 |
| receptor ligand activity (GO:0048018) | 0.321 | 0.354 | 2.7 | 3.0 |
| calcium ion binding (GO:0005509) | 0.355 | 0.382 | 2.4 | 2.4 |
| purine ribonucleoside triphosphate binding (GO:0035639) | 0.441 | 0.463 | 1.8 | 1.4 |
| protein homodimerization activity (GO:0042803) | 0.554 | 0.568 | 1.3 | 0.7 |

| | | | | |
|---|---|---|---|---|
| DNA binding (GO:0003677) | 0.645 | 0.645 | 1.0 | 0.4 |

**Supplementary Table 3:** Growth conditions for cell lines used in this study

| *Cell Line* | *Plating density* | *Growth media* | *Growth conditions* | *Subtype* | *Notes* |
|---|---|---|---|---|---|
| 184A1 | 1250 | MEBM (CC-3150) + 1% FBS + 1% P/S | 37˚C, 5% $CO_2$ | NM, basal[1] | |
| AU565 | 1250 | RPMI-1640 + 10% FBS + 1% P/S | 37˚C, 5% $CO_2$ | $HER2^{amp}$, luminal[2] | INPP4B-low[3] |
| BT20 | 1500 | EMEM + 10% FBS + 1% P/S | 37˚C, 5% $CO_2$ | TNBC, basal[2] | PIK3CA-kin[4] |
| BT474 | 1750 | RPMI-1640 + 10% FBS + 1% P/S | 37˚C, 5% $CO_2$ | $HER2^{amp}$, luminal[2] | |
| BT549 | 1000 | RPMI-1640 + 10% FBS + 1% P/S, 1 ug/ml IN | 37˚C, 5% $CO_2$ | TNBC, basal[2] | PTEN-low[3], pRb-def[5] |
| CAL120 | 750 | DMEM + 10% FBS + 1% P/S | 37˚C, 5% $CO_2$ | TNBC, basal[6] | |
| CAL51 | 1000 | DMEM + 20% FBS + 1% P/S | 37˚C, 5% $CO_2$ | TNBC, basal[6] | PIK3CA-hel[4] |
| CAL851 | 1500 | DMEM + 10% FBS + 1% P/S | 37˚C, 5% $CO_2$ | TNBC, basal[6] | pRb-def[5] |
| CAMA1 | 1750 | EMEM + 10% FBS + 1% P/S | 37˚C, 5% $CO_2$ | $HR^+$, luminal[2] | |
| EFM19 | 1750 | RPMI-1640 + 10% FBS + 1% P/S | 37˚C, 5% $CO_2$ | $HR^+$, luminal[6] | PIK3CA-kin[4] |
| EVSAT | 1000 | EMEM + 10% FBS + 2 mM L-glutamine, 1% P/S | 37˚C, 5% $CO_2$ | $HR^+$, luminal[6] | |
| HCC1143 | 750 | RPMI-1640 + 10% FBS + 1% P/S | 37˚C, 5% $CO_2$ | TNBC, basal[2] | |
| HCC1187 | 2500 | RPMI-1640 + 10% FBS + 1% P/S | 37˚C, 5% $CO_2$ | TNBC, basal[2] | |
| HCC1395 | 1500 | RPMI-1640 + 10% FBS + 1% P/S | 37˚C, 5% $CO_2$ | TNBC, basal | PTEN-low[3] |
| HCC1419 | 1000 | RPMI-1640 + 10% FBS + 1% P/S | 37˚C, 5% $CO_2$ | $HER2^{amp}$, luminal[1] | |
| HCC1428 | 1000 | RPMI-1640 + 10% FBS + 1% P/S | 37˚C, 5% $CO_2$ | $HR^+$, luminal[2] | |

| | | | | | |
|---|---|---|---|---|---|
| HCC1500 | 2000 | RPMI-1640 + 10% FBS + 1% P/S | 37˚C, 5% $CO_2$ | $HR^+$, luminal[1] | |
| HCC1569 | 1750 | RPMI-1640 + 10% FBS + 1% P/S | 37˚C, 5% $CO_2$ | $HER2^{amp}$, basal[2] | PTEN-low[3], pRb-def[5] |
| HCC1806 | 1000 | RPMI-1640 + 10% FBS + 1% P/S | 37˚C, 5% $CO_2$ | TNBC, basal[6] | INPP4B-low[3] |
| HCC1937 | 1500 | RPMI-1640 + 10% FBS + 1% P/S | 37˚C, 5% $CO_2$ | TNBC, basal[2] | PTEN-low[3], pRb-def[5] |
| HCC1954 | 750 | RPMI-1640 + 10% FBS + 1% P/S | 37˚C, 5% $CO_2$ | $HER2^{amp}$, basal[2] | PIK3CA-kin[4] |
| HCC202 | 2250 | RPMI-1640 + 10% FBS + 1% P/S | 37˚C, 5% $CO_2$ | $HER2^{amp}$, luminal[2] | PIK3CA-hel[4] |
| HCC38 | 1000 | RPMI-1640 + 10% FBS + 1% P/S | 37˚C, 5% $CO_2$ | TNBC, basal[2] | PTEN-low[3] |
| HCC70 | 2000 | RPMI-1640 + 10% FBS + 1% P/S | 37˚C, 5% $CO_2$ | TNBC, basal[2] | PTEN-low[3], pRb-def[5] |
| HS578T | 500 | DMEM + 10% FBS + 1% P/S | 37˚C, 5% $CO_2$ | TNBC, basal[2] | INPP4B-low[3] |
| HTERTHME1 | 1000 | MEMB + Lonza CC-3150 kit | 37˚C, 5% $CO_2$ | NM, basal[1] | |
| MCF10A | 1000 | DMEM/F12 (1:1) + 5% HS + 1% P/S, 20 ng/ml EGF, 0.5 mg/ml HC, 10 µg/ml IN, 100 ng/ml CT | 37˚C, 5% $CO_2$ | NM, basal[2] | |
| MCF12A | 400 | DMEM/F12 + 5% HS + 1% P/S, 20 ng/ml EGF, 0.5 mg/ml HC, 10 µg/ml IN, 100 ng/ml CT | 37˚C, 5% $CO_2$ | NM, basal[2] | INPP4B-low[3] |
| MCF7 | 2000 | DMEM + 10% FBS + 1% P/S | 37˚C, 5% $CO_2$ | $HR^+$, luminal[2] | PIK3CA-hel[4] |
| MDAMB157 | 1000 | L-15 + 10% FBS + 1% P/S | 37˚C, 0% $CO_2$ | TNBC, basal[2] | INPP4B-low[3] |
| MDAMB175VII | 2500 | L-15 + 10% FBS + 2mM L-glutamine, 1% P/S | 37˚C, 0% $CO_2$ | $HR^+$, luminal[2] | |
| MDAMB231 | 1000 | DMEM + 10% FBS + 1% P/S | 37˚C, 5% $CO_2$ | TNBC, basal[2] | |

| | | | | | |
|---|---|---|---|---|---|
| MDAMB330 | 2500 | L-15 + 20% FBS + 2mM L-glutamine,1% P/S + 30 ng/ ml EGF + 0.016 mg/ml IN + 2mM glutathione | 37˚C, 0% $CO_2$ | $HER2^{amp}$, luminal[6] | |
| MDAMB361 | 1500 | L-15 + 20% FBS + 1% P/S | 37˚C, 0% $CO_2$ | $HER2^{amp}$, luminal[2] | PIK3CA-hel[4] |
| MDAMB415 | 2500 | L-15 + 15% FBS + 2mM L-glutamine,1% P/S, 10 µg/ml IN | 37˚C, 0% $CO_2$ | $HR^+$, luminal[2] | |
| MDAMB436 | 1000 | L-15 + 10% FBS + 1% P/S, 10 µg/ml IN | 37˚C, 0% $CO_2$ | TNBC, basal[2] | PTEN-low[3], pRb-def[5] |
| MDAMB453 | 1500 | L-15 + 10% FBS + 1% P/S | 37˚C, 0% $CO_2$ | TNBC/$HER2^{amp}$, luminal[2] | PIK3CA-kin[4] |
| MDAMB468 | 1000 | L-15 + 10% FBS + 1% P/S | 37˚C, 0% $CO_2$ | TNBC, basal[2] | PIK3CA-hel[4], pRb-def[5] |
| MGH312 | 2000 | RPMI + 10% FBS + 1% P/S | 37˚C, 5% $CO_2$ | $HR^+$, luminal[5] | pRb-def[5] |
| PDX1206 | 2000 | DMEM/F12 (3:1) + 7.5% FBS + 1% P/S, 0.125 ng/ml EGF, 25 ng/ml HC, 5 µg/ml IN, 8.6 ng/ml CT, 5 uM Y-27632 | 37˚C, 5% $CO_2$ | TNBC, basal | |
| PDX1258 | 1500 | DMEM/F12 (3:1) + 7.5% FBS + 1% P/S, 0.125 ng/ml EGF, 25 ng/ml HC, 5 µg/ml IN, 8.6 ng/ml CT, 5 uM Y-27632 | 37˚C, 5% $CO_2$ | TNBC, basal | pRb-def[5] |
| PDX1328 | 1500 | DMEM/F12 (3:1) + 7.5% FBS + 1% P/S, 0.125 ng/ml EGF, 25 ng/ml HC, 5 µg/ml IN, 8.6 ng/ml CT, 5 uM Y-27632 | 37˚C, 5% $CO_2$ | TNBC, basal | |

| | | | | | |
|---|---|---|---|---|---|
| PDXHCI002 | 1500 | DMEM/F12 (3:1) + 7.5% FBS + 1% P/S, 0.125 ng/ml EGF, 25 ng/ml HC, 5 µg/ml IN, 8.6 ng/ml CT, 5 uM Y-27632 | 37˚C, 5% $CO_2$ | TNBC, basal[7] | |
| SKBR3 | 1500 | McCoy's + 10% FBS + 1% P/S | 37˚C, 5% $CO_2$ | $HER2^{amp}$, luminal[2] | INPP4B-low[3] |
| SUM1315 | 1000 | F-12 + 5% FBS + 1% P/S, 10 ng/ml EGF, 5 µg/ml IN, 10 mM HEPES | 37˚C, 5% $CO_2$ | TNBC, basal[2] | INPP4B-low[3] |
| SUM149 | 1000 | F-12 + 5% FBS + 1% P/S, 1 µg/ml HC, 5 µg/ml IN, 10 mM HEPES | 37˚C, 5% $CO_2$ | TNBC, basal[2] | PTEN-low[3] |
| SUM159 | 1000 | F-12 + 5% FBS + 1% P/S, 1 µg/ml HC, 5 µg/ml IN, 10 mM HEPES | 37˚C, 5% $CO_2$ | TNBC, basal[2] | PIK3CA-kin[8] |
| SUM185PE | 2500 | Ham's F12 + 5% FBS + 1% P/S + 5 µg/ml BI + 1 µg/ml HC + 10 mM HEPES | 37˚C, 5% $CO_2$ | TNBC, luminal[2] | PIK3CA-kin[8] |
| SUM190PT | 2000 | Ham's F12 + 1% FBS + 1% P/S, 5 µg/ml BI, 1 µg/ml HC, 10 mM Hepes, 5 mM ethanolamine, 5 µg/ml transferrin, 10 nM triiodo thyronine, 50 nM sodium selenite, 1 g/l BSA | 37˚C, 5% $CO_2$ | $HER2^{amp}$, basal[2] | PIK3CA-kin[8] |
| SUM229PE | 1000 | Ham's F12 + 5% FBS + 1% P/S + 5 µg/ml BI + 1 µg/ml HC + 10 mM HEPES | 37˚C, 5% $CO_2$ | TNBC, basal[6] | |
| SUM44PE | 1750 | Ham's F12 + 1% FBS + 1% P/S, 5 µg/ml BI, 1 µg/ml HC, 10 | 37˚C, 5% $CO_2$ | $HR^+$, luminal[2] | |

| | | | | | |
|---|---|---|---|---|---|
| | | mM HEPES, 5 mM ethanolamine, 5 µg/ml transferrin, 10 nM triiodo thyronine, 50 nM sodium selenite, 1 g/l BSA | | | |
| SUM52PE | 1500 | Ham's F12 + 5% FBS + 1% P/S + 5 µg/ml BI + 1 µg/ml HC + 10 mM HEPES | 37˚C, 5% $CO_2$ | $HR^+$, luminal[2] | INPP4B-low[3] |
| T47D | 1000 | RPMI-1640 + 10% FBS + 1% P/S, 1 µg/ml IN | 37˚C, 5% $CO_2$ | $HR^+$, luminal[2] | PIK3CA-kin[4] |
| UACC812 | 2500 | L-15 + 20% FBS + 2mM L-glutamine, 1% P/S, 20 ng/ml EGF | 37˚C, 0% $CO_2$ | $HER2^{amp}$, luminal[2] | |
| UACC893 | 2500 | L-15 + 10% FBS + 1% P/S | 37˚C, 0% $CO_2$ | $HER2^{amp}$, luminal[6] | PIK3CA-kin[4] |
| ZR-75-1 | 1500 | RPMI-1640 + 10% FBS + 1% P/S | 37˚C, 5% $CO_2$ | $HR^+$, luminal[2] | PTEN-low[3] |
| ZR-75-30 | 1750 | RPMI-1640 + 10% FBS + 1% P/S | 37˚C, 5% $CO_2$ | $HER2^{amp}$, luminal[2] | |

BI: Bovine insulin, CT: Cholera toxin, EGF: Epidermal growth factor, FBS: Fetal bovine serum, HC: Hydrocortisone, HS: Horse serum, IN: Insulin, P/S: Penicillin/Streptomycin, pRb-def: pRb deficient, PIK3CA-hel: PIK3CA helical domain mutation, PIK3CA-kin: PIK3CA kinase domain mutation. Note: in cases when multiple PI3K pathway activation changes were present in a cell line, we prioritized PIK3CA mutations, followed by low PTEN and then low INPP4B for the purposes of plotting Fig. 4.

**Supplementary Table 4:** Nominal targets and pathways inhibited by drugs used in this study

| ***Agent*** | ***Synonyms*** | ***Nominal target*** | ***Nominal pathway*** | ***HMSLID*** |
|---|---|---|---|---|
| A-1210477 | | Mcl-1 | Bcl2 family | 10478-102 |
| Abemaciclib | LY2835219, Verzenio | CDK4/6 | CDK - cell cycle | 10390-103 |
| ABT-737 | | Bcl2/XL | Bcl2 family | 10179-101 |
| Alpelisib | BYL719, Piqray | PI3Ka | PI3K/AKT/mTOR | 10233-101 |
| Alvocidib | Flavopiridol, L86-8275, HMR-1275 | pan-CDK | CDK | 10011-101 |

| | | | | |
|---|---|---|---|---|
| Adavosertib | AZD1775, MK1775 | WEE1 | Cell cycle | 10152-101 |
| Vistusertib | AZD2014 | mTor | PI3K/AKT/mTOR | 10437-101 |
| Capivasertib | AZD5363 | Akt | PI3K/AKT/mTOR | 10370-101 |
| Ceralasertib | AZD6738 | ATR | DNA damage | 10677-101 |
| AZD7762 | AZD7762 | CHK1/2 | Cell cycle | 10006-101 |
| BJP-6-5-3 | | Pin 1 | Other | N/A |
| Bleomycin | | Radiomimetic | DNA damage | 10482-114 |
| BMS-265246 | | CDK1/2 | CDK - cell cycle | 10680-101 |
| BSJ-01-175 | | CDK12/13 | CDK | N/A |
| BSJ-03-123 | | CDK6 | CDK - cell cycle | N/A |
| BSJ-03-124 | | CDK4/6 degron | CDK - cell cycle | N/A |
| Buparlisib | BKM120 | pan PI3K | PI3K/AKT/mTOR | 10147-101 |
| Ulixertinib | BVD523, VRT752271 | ERK1/2 | MAPK/ERK | 10665-101 |
| Cabozantinib | BMS-907351, XL184, Cabometyx, Cometriq | VEGFR2/MET | RTK | 10194-106 |
| Cediranib | AZD2171 | VEGFR/cKIT | RTK | 10444-101 |
| Ceritinib | LDK378, Zykadia | ALK | RTK | 10394-101 |
| CFI-400945 | | PLK4 | Cell cycle | N/A |
| Cisplatin | CDDP, cis-Diaminodichloroplatinum | DNA | DNA damage | 10245-999 |
| Dasatinib | BMS-354825, Sprycel | BCR/ABL | MAPK/ERK | 10020-999 |
| Dinaciclib | SCH727965 | pan CDK | CDK | 10287-101 |
| Doxorubicin | Hydroxydaunorubicin | DNA | DNA damage | 10248-102 |
| E17 | | CDK12/13 | CDK | N/A |
| Etoposide | VP-16, VP-16-213, Etopophos, Toposar | Topoisomerase II | DNA damage | 10250-101 |
| Everolimus | RAD001, SDZ-RAD, Zortress, Afinitor Disperz, Afinitor | mTOR1 | PI3K/AKT/mTOR | 10235-101 |
| FMF-03-145-1 | | pan-Aurora | Cell cycle | N/A |

| | | | | |
|---|---|---|---|---|
| FMF-03-146-1 | | DCLK1/2 | Other | N/A |
| FMF-04-107-2 | | CDK14 | CDK | N/A |
| FMF-04-112-1 | | DCLK1/2 | Other | N/A |
| GSK2334470 | | PDK1 | PI3K/AKT/mTOR | 10221-101 |
| Sapanisertib | INK128, MLN0128, TAK228 | mTORC1/2 | PI3K/AKT/mTOR | 10205-101 |
| Ipatasertib | GDC0068, RG7440 | AKT | PI3K/AKT/mTOR | 10338-101 |
| Luminespib | AUY922, VER-52296 | HSP90 | Other | 10161-101 |
| Prexasertib | LY2606368 | Chk1 | Cell cycle | N/A |
| Samotolisib | LY3023414 | PI3K/mTor/DNA-PK | PI3K/AKT/mTOR | 10676-101 |
| MFH-2-90 | | CDK12/13 | CDK | N/A |
| Neratinib | HKI272, Nerlynx | EGFR/HER2 | RTK | 10018-999 |
| Olaparib | AZD2281, KU0059436, Lynparza | PARP | DNA damage | 10144-101 |
| Paclitaxel | Taxol, Abraxane | Tubulin | Chemotherapy | 10102-999 |
| Palbociclib | PD0332991, Ibrance | CDK4/6 | CDK - cell cycle | 10071-102 |
| PF-4708671 | | p70S6K | PI3K/AKT/mTOR | 10404-101 |
| Pictilisib | GDC0941 | pan PI3K | PI3K/AKT/mTOR | 10047-101 |
| Pin1-3 | | Pin 1 | Other | N/A |
| R0-3306 | | CDK1 | CDK - cell cycle | 10104-101 |
| Ribociclib | LEE011, Kisqali | CDK4/6 | CDK - cell cycle | 10395-101 |
| Rucaparib | AG014699, PF-01367338, Rubraca | PARP | DNA damage | 10304-105 |
| Saracatinib | AZD0530 | SRC | MAPK/ERK | 10032-101 |
| Senexin b | | CDK8/19 | CDK | 10679-101 |
| SHP099 | | SHP2 | MAPK/ERK | 10664-102 |
| Taselisib | GDC0032, RG-7604 | PI3Ka, g, d | PI3K/AKT/mTOR | 10445-101 |
| TGX221 | | PI3Kb | PI3K/AKT/mTOR | 10171-101 |
| THZ-P1-2 | | PIP4K | PI3K/AKT/mTOR | N/A |
| THZ-P1-2R | | PIP4K | PI3K/AKT/mTOR | N/A |
| THZ1 | | CDK7/12/13 | CDK | 10429-101 |
| THZ531 | | CDK12/13 | CDK | 11996-999 |
| Tivantinib | ARQ197 | MET | RTK | 10131-101 |

| | | | | |
|---|---|---|---|---|
| Topotecan | SKF104864A, NSC609669, Hycamtin, Potactasol | Topoisomerase I | DNA damage | 10278-102 |
| Torin 2 | | mTOR/PIKKs | PI3K/AKT/mTOR | N/A |
| Trametinib | GSK1120212, JTP-74057, Mekinist | MEK | MAPK/ERK | 10142-101 |
| Volasertib | BI6727 | PLK | Cell cycle | 10387-101 |
| Vorinostat | SAHA, Zolinza | HDAC | Other | 10282-101 |
| YKL-5-124 | | CDK7 | CDK | N/A |
| ZZ1-33B | | CDK9 degron | CDK | N/A |

**Supplementary Table 5:** Metadata for antibodies used in cyclic immunofluorescence shown in Fig. 3a-b, Hoechst was included in each cycle.

| ***Cycle*** | ***Antigen*** | ***Dilution used*** | ***Clone*** | ***Fluor*** | ***Vendor*** | ***Cat #*** | ***Lot #*** | ***RRID*** |
|---|---|---|---|---|---|---|---|---|
| 1 | pH3 | 1:1000 | D2C8 | 488 | CST | 3465S | 14 | AB_10695860 |
| 1 | pRb | 1:500 | D20B12 | 555 | CST | 8957S | 5 | AB_2728827 |
| 1 | b-catenin | 1:300 | L54E2 | 647 | CST | 4627S | 5 | AB_10691326 |
| 2 | PCNA | 1:300 | PC10 | 488 | CST | 8580S | 3 | AB_11178664 |
| 2 | ki67 | 1:300 | 20Raj1 | 570 | Thermo | 41-5699-82 | 1956875 | AB_11220278 |
| 2 | p21 | 1:300 | 12D1 | 647 | CST | 8587S | 6 | AB_10892861 |
| 3 | Cyclin D1 | 1:100 | EPR2241 | 488 | Abcam | 190194 | GR3282485-2 | AB_2728784 |
| 3 | b-actin | 1:500 | 13E5 | 555 | CST | 8046S | 3 | AB_11179208 |
| 3 | g-H2AX | 1:300 | 20E3 | 647 | CST | 9720S | 19 | AB_10692910 |
